# Structural insights into context-specific inhibition of bacterial translation by macrolides

**DOI:** 10.1101/2025.06.03.657637

**Authors:** Egor A. Syroegin, Elena V. Aleksandrova, Artem A. Kruglov, Madhura N. Paranjpe, Maxim S. Svetlov, Yury S. Polikanov

**Affiliations:** Department of Biological Sciences, University of Illinois at Chicago, Chicago, IL 60607, USA; Department of Pharmaceutical Sciences, University of Illinois at Chicago, Chicago, IL 60607, USA; Center for Biomolecular Sciences, University of Illinois at Chicago, Chicago, IL 60607, USA

**Keywords:** Macrolides, erythromycin, telithromycin, antibiotic, peptidyl-tRNA, 70S ribosome, X-ray structure, peptidyl transferase center, nascent peptide exit tunnel, context-specificity of drug action, inhibition of protein biosynthesis

## Abstract

The ribosome’s peptidyl transferase center (PTC) catalyzes peptide bond formation during protein synthesis and is targeted by many antibiotic classes. Remarkably, macrolides that bind in the peptide exit tunnel some ∼10Å away from the PTC also remotely inhibit PTC and cause translational arrest depending on the synthesized polypeptide sequence. The Arg/Lys-X-Arg/Lys (also known as +X+) motif is particularly susceptible to this inhibition, as peptidyl-tRNA carrying nascent peptide with penultimate arginine or lysine residue fails to react with aminoacyl-tRNA carrying the same amino acids in the presence of macrolides. While structural studies of macrolide-bound ribosomes have shed light on the context-specific nature of this inhibition, the precise roles of the drug, ribosome, and tRNA in modulating PTC activity remain unclear. In this study, we present a detailed structural analysis of ribosome-nascent chain complexes (RNCs) that represent either arrested or non-arrested states, containing various combinations of peptidyl- and aminoacyl-tRNAs, with or without macrolides. Our findings reveal a dynamic interaction between the ribosome-bound drug, the nascent peptide, and the incoming amino acid, which collectively modulates PTC function. This lays the foundation for designing antibiotics that can overcome drug resistance by preventing the induction of inducible *erm* genes in pathogens.

## INTRODUCTION

Protein biosynthesis, also known as translation, is a complex multi-step process essential to every cell and is catalyzed by the ribosome. During this process, the catalytic peptidyl transferase center (PTC) located at the heart of the large ribosomal subunit polymerizes amino acids into polypeptides in the order specified by the nucleotide sequences of the translated mRNAs. Newly-made proteins emerge from the ribosome through the 100Å-long nascent peptide exit tunnel (NPET) that spans the body of the large subunit, beginning at the PTC and ending on the opposite side of the subunit. Remarkably, the PTC effectively catalyzes transpeptidation reactions between a wide variety of peptidyl- and aminoacyl-tRNAs, with only a few “problematic” substrate combinations that require the assistance of auxiliary translation factors^1,2^.

The ribosome is one of the major targets in bacterial cells for antimicrobial drugs that are indispensable as therapeutic agents and tools for basic research^3,4^. Macrolides and their newest generation, ketolides, represent a clinically significant family of antibiotics active against a broad spectrum of Gram-positive bacteria^5^. These drugs bind and act in the NPET of the bacterial ribosome, close to the PTC. It has been thought for years that macrolides inhibit translation by simply clogging the NPET, thereby blocking the passage of all newly made polypeptides once they grow to the size of 3-10 amino acids^6–10^. However, in the past decade, this simplistic view of macrolides acting like mere plugs of the NPET has been significantly transformed. One striking observation that the initial model of macrolide action could not easily explain was that protein synthesis is not completely inhibited and ribosome can still efficiently synthesize a subset of full-length proteins *in vivo* even at concentrations of macrolides exceeding by many fold the minimal inhibitory concentration (MIC) that prevents cell growth^11,12^. Remarkably, rather than equally curtailing the synthesis of all proteins, macrolides abolish the production of a number of polypeptides, whereas some others continue to be translated at levels comparable to those in untreated cells^11,12^. Subsequent re-examination of the structures of macrolides in complex with the ribosome showed that the opening of the NPET occupied by a macrolide (such as erythromycin, ERY) appears to be sufficiently wide for at least some nascent peptides to squeeze through^13,14^. Later, the cryo-EM structures of the ERY-bound ribosome carrying a nascent peptide demonstrated that a protein chain can be comfortably accommodated in the NPET together with the antibiotic^15–17^.

The most remarkable finding from the ribosome profiling data is that translation of many genes is arrested at relatively few well-defined sites in the presence of macrolides^18,19^. Analysis of these sites showed that they are defined by specific sequence signatures within the nascent peptide chain, Macrolide Arrest Motifs (MAMs)^18,19^. One of the most prevalent MAMs, accounting for ∼80% of the strongest ribosome stalling sites in the presence of macrolides, conforms to the consensus sequence Lys/Arg-X-Lys/Arg, in which the first two residues belong to the C-terminal part of the P-site peptidyl-tRNA, and the third residue is the incoming aminoacyl-tRNA (aa-tRNA) (**Figure S1**). In other words, the A-site of the macrolide-arrested ribosomes discriminates against incoming Arg/Lys aa-tRNAs if Arg/Lys appear in the penultimate (-1) position of the nascent chain. Since the positive charges of the -1 amino acid residue of the P-site donor and the A-site aminoacyl acceptor are the key determinants of the problematic nature of this MAM, it was also referred to as +X+ motif^20^. Importantly, chemically diverse macrolides, including prototypic ERY^18,20,21^, telithromycin (TEL)^18,21,22^, solithromycin^20^, and clinically widely used azithromycin (AZI)^19,20^ induce ribosome stalling at this motif both *in vivo* and *in vitro*, suggesting that universal chemical features shared by all macrolides cause stalling.

In the macrolide-stalled ribosome, the arrest motif is located at the PTC, but it is considered too distant to directly interact with the macrolide bound in the NPET (**Figure S1**), suggesting that macrolides alter the selectivity of the A site indirectly, and allosterically predispose PTC to translation arrest. Therefore, MAMs cause problems not because the growing peptide is trapped in the drug-obstructed tunnel but because the macrolide-bound ribosome is unable to catalyze peptide bond formation between the last two residues of a MAM, such as +X+ motif^5,13,15,16,20,23^. As a result, macrolides selectively abolish the production of a subset of cellular proteins having specific amino acid sequences^11^. Rather than being simple tunnel plugs, macrolide antibiotics emerge as context-specific inhibitors of peptide bond formation – a feature that unites them with PTC-targeting antibiotics, such as chloramphenicol (CHL).

Importantly, MAMs, such as the +X+, are found in many regulatory leader upstream ORFs (uORFs) that control the expression of clinically important macrolide-resistance genes, such as *erm*^20,24,25^, *mef*^24,26^, or *msr*^24,26^. Since the expression of these genes can significantly reduce cell fitness, their activation often occurs only when bacteria are under antibiotic stress. In the absence of the drug, ribosomes effectively synthesize leader peptide from a bicistronic mRNA, but the translation of the downstream resistance gene is suppressed due to the mRNA secondary structure hindering its ribosome-binding site^24,27^. However, in the presence of an inducing macrolide, ribosomes stall at the +X+ motif of the leader peptide sequence, re-arranging the downstream mRNA structure, exposing the ribosome-binding site, and, thereby, permitting translation of the resistance gene.

A well-known example of a leader uORF containing the +X+ motif is ErmDL (MTHSM**R**L**R**IFPTL), a peptide that regulates the expression of the ErmD methyltransferase, which confers resistance to macrolides^28^. Macrolides stall translation of ErmDL peptide when ribosome contains P-site fMTHSMRL-peptidyl-tRNA^Leu^ and A-site arginyl-tRNA^Arg^. Interestingly, even when truncated to just the three C-terminal residues (MRL), this peptide is still capable of stalling the ribosome in response to macrolide binding. Recent structural analysis of ribosome-nascent chain complexes (RNCs) containing the ErmDL peptide has shown that macrolide ERY (or the ketolide TEL) induces a specific conformation of the nascent peptide, where the penultimate arginine residue extends into the A-site cleft^21^. In this drug-induced conformation, the penultimate arginine clashes directly with the incoming arginyl-tRNA, likely preventing its productive binding to the translating ribosome^21^. While this work suggested that the interplay between the drug, the ribosome, and the ErmDL peptide mediates translational arrest, it remains unclear

### what are the specific roles that the antibiotic, the residues of the peptide, or the amino acid of the incoming A-site tRNA play in the translational arrest?

The primary challenge in resolving these questions lies in the absence of reference structures representing non-arrested RNCs, such as those without bound macrolide, with a mutated arrest peptide sequence or with a non-Arg/Lys A-site amino acid. Additionally, while ErmDL provides a well-studied example of macrolide-induced stalling, it remains uncertain how generalizable its mechanism is across all +X+ arrest sites.

In this work, by utilizing stably-linked hydrolysis-resistant aminoacyl- and peptidyl-tRNA analogs, we report a series of high-resolution X-ray crystal structures of the *Thermus thermophilus* (*Tth*) 70S ribosome with or without bound macrolides, as well as carrying active (i.e., stalling-prone) or inactive combinations of aminoacyl- and peptidyl-tRNAs in the A and P sites, respectively. The key insight from our study is the identification of a previously unseen conformation of the lysyl-tRNA in the A site, in which the side chain of the incoming lysine cannot fully accommodate into the A-site cleft, causing lysine’s α-amino group to be positioned prohibitively far from the carbonyl group of the peptidyl-tRNA making peptide bond formation unattainable. We reveal that the binding of macrolides in the NPET disrupts the accommodation of some aa-tRNAs in the A-site, particularly when the incoming tRNA carries large and flexible amino acids such as arginine or lysine. In the presence of macrolides, the penultimate arginine residue of the nascent peptide chain adopts a conformation that obstructs the A-site, preventing proper tRNA binding and peptide bond formation. This structural disruption is specific to sequences containing the +X+ motif, where the macrolide delicately modulates the interaction between the ribosome, nascent peptide, and incoming tRNA. Our findings provide key insights into how the binding of small molecules, such as macrolides, within the ribosomal tunnel can modulate bacterial translation depending on the sequence of a synthesized peptide. This new perspective fundamentally extends our understanding of how small ligands convert ribosome from a universal protein synthesis machine to a selective producer of specific polypeptides.

## RESULTS AND DISCUSSION

### Short ErmDL sequence facilitates analysis of arrested and non-arrested RNCs

The main goal of this study is to provide the structural bases for the context-specific action of macrolides on +X+ motifs – the most common type of MAMs in bacteria^18,19^. Specifically, we aimed to understand the roles of the positively charged amino acid residues at the penultimate (-1) position of the growing polypeptide chain and the incoming A-site aa-tRNA, which are critical for macrolide activity^20^. Previous research has shown that the N-terminal part of the ErmDL peptide can stall the ribosome in a (+X+)-independent manner^21^, complicating the interpretation of how the +X+ motif itself contributes to macrolide-induced translational arrest. To address this, we chose the MRLR sequence, which is the shortest known functional +X+ motif, allowing us to eliminate the influence of N-terminal residues and study the pure +X+ motif. Moreover, the reported structures of macrolide-stalled RNCs translating the ErmDL sequence^21^ cannot explain how a short MRL-tripeptidyl-tRNA can mediate ribosome stalling, given that its residues might be too distant to directly interact with the macrolide molecule^20^.

Determining the precise orientations of essential components involved in macrolide-induced translation arrest is crucial for understanding why PTC fails to catalyze peptide bond formation during stalling events. To dissect the individual roles of these components, structural comparisons between arrested and non-arrested RNCs are necessary (e.g., with vs. without antibiotic, WT vs. mutant MRL sequence, Lys/Arg-tRNA vs. other aa-tRNAs). Previously, such analyses were impossible due to the reliance on cryo-EM structures of only the arrested RNCs, which inherently captured an inactive PTC state^15–17,21^. The *in vitro* translation-based approach used in those studies relies on ribosome stalling in the presence of a macrolide and an arrest peptide, making it unsuitable for generating corresponding non-arrested RNCs. An alternative method involves reconstituting arrested and non-arrested RNCs from purified, non-reactive components^29–31^. Therefore, to uncover the mechanism of macrolide-induced context-specific PTC inhibition, we employed our recently developed chemoenzymatic approach based on native chemical ligation (NCL)^29^, enabling the preparation of non-hydrolyzable full-length peptidyl-tRNA carrying either formyl-Met-Arg-Cys (fMRC) or formyl-Met-Ala-Cys (fMAC) tripeptide moieties. The flexibility of the +X+ motif sequence, where the identity of the “X” residue is non-essential, is crucial for this study, as it allows the incorporation of cysteine at the peptide’s C-terminus, a requirement dictated by NCL reaction chemistry.

To ensure that the peptide sequences of the peptidyl-tRNAs and their combinations with various aa-tRNAs indeed support or counteract macrolide action during translation, we carried out primer extension inhibition assays (toe-printing), which allows detection of the drug-induced ribosome stalling site(s) along mRNAs with single-codon accuracy^32^. As expected, the addition of ERY to the cell-free translation system programmed with the MRLR mRNA resulted in ribosome stalling at the Leu3 codon of this ORF, when arginine residues appeared in the penultimate (-1) position of the growing polypeptide chain and also as the incoming (+1) amino acid (**Figure S2**, lane 2, red arrowhead). Importantly, alteration of the +1 arginine to lysine (**Figure S2**, lane 4) or the C-terminal leucine to cysteine (**Figure S2**, lane 6) preserves ribosome stalling at the same site. In contrast, as expected, the replacement of the -1 arginine to alanine or the +1 arginine to phenylalanine or proline abolished ERY-dependent ribosome stalling (**Figure S2**, lanes 8, 10, and 12, red arrowhead), which is fully consistent with the reported context specificity of ERY action^18–20^.

### Macrolides prevent productive accommodation of lysyl-tRNA into the A site

Using purified components, we assembled WT *Thermus thermophilus (Tth)* 70S ribosome complexes recapitulating macrolide-arrested RNCs containing non-hydrolyzable fMRC-peptidyl-tRNAi^Met^ in the P site, lysyl-tRNA^Lys^ in the A site, and either macrolide ERY or ketolide TEL, crystallized them, and determined their X-ray structures at 2.65 and 2.8 Å resolution, respectively (**Figure 1; Figure S3; Table S1**). In the resulting structures, both drugs bind in the canonical macrolide pocket within the NPET that is formed by nucleotides A2058 and A2059 of the 23S rRNA (**Figure 1A, B; Figure S3A, B**) and adopt conformations indistinguishable from those previously observed in the context of non-stalled *Tth* 70S ribosome (**Figure S4A, B**)^33^ or stalled *E. coli* 70S RNCs (**Figure S4C, D**)^21^. In both structures, the desosamine moiety of a macrolide (ERY or TEL) forms two hydrogen bonds (H-bonds) crucial for binding^33^: one between the hydroxyl group of the desosamine and the N1 atom of the nucleotide A2058, and the second one between the dimethylamino group of desosamine sugar and the N6 atom of A2058 via a water molecule that is also coordinated by the phosphate of G2505 (**Figure 1C, D**). While the observed electron density for the body, and especially the elbow region of the A-site tRNA was relatively weak, the density for the anticodon loop and the CCA-end were well-resolved (**Figure 1C**). Importantly, the obtained electron density maps allowed confident modeling of the lysine residue of the aminoacylated lysyl-tRNA^Lys^ in the A site and peptide moiety of the fMRC-tripeptidyl-tRNA in the P site (**Figure 1C, D**).

**Figure 1.**
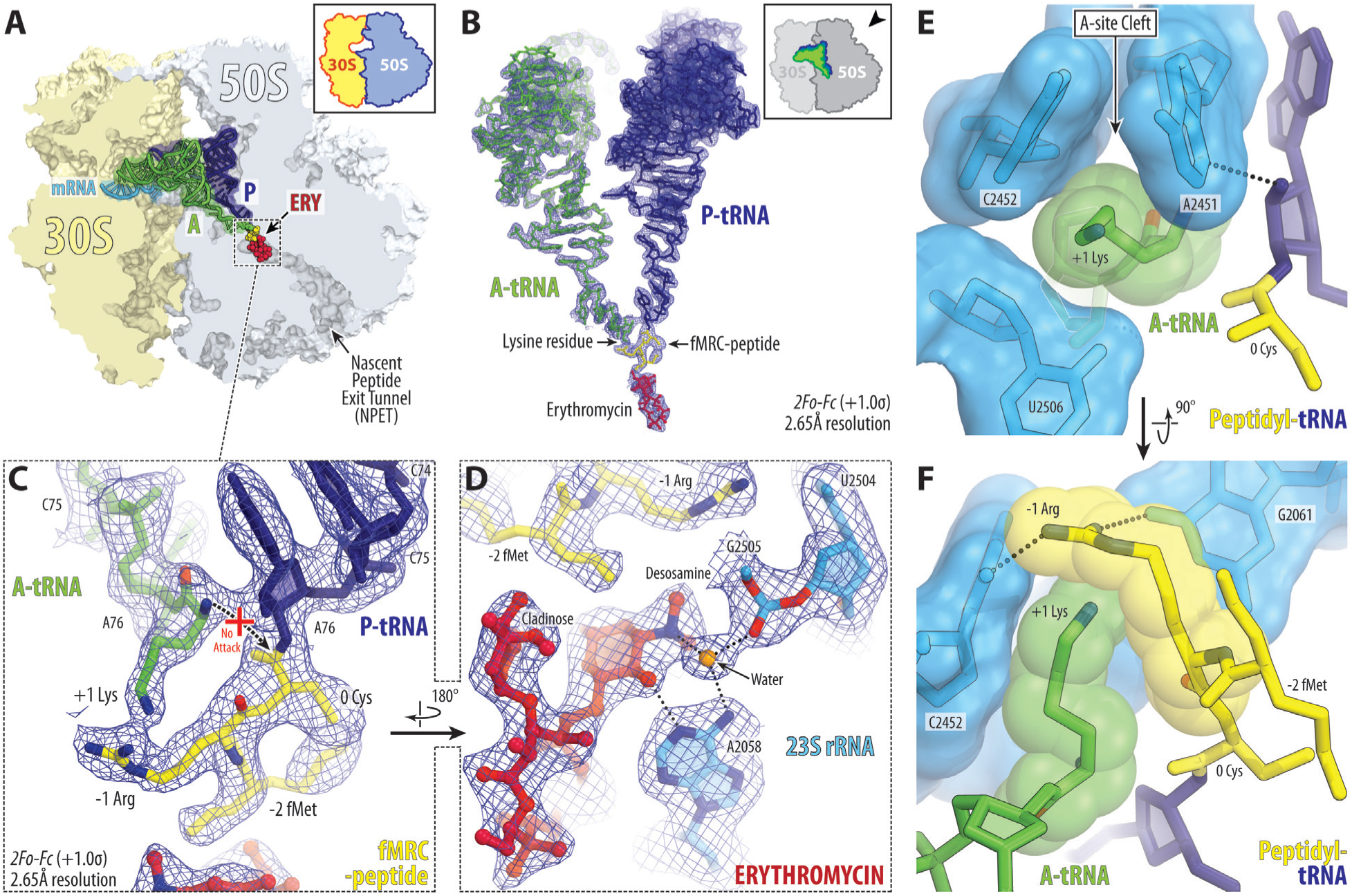
Structure of the erythromycin-arrested RNC assembled from purified components. (**A**) Overview of the structure of the *T. thermophilus* 70S ribosome in complex with erythromycin (ERY, red) and also containing non-hydrolyzable lysyl-tRNA^Lys^ and fMRC-peptidyl-tRNAi^Met^ in the A and P sites, respectively. The 30S subunit is shown in light yellow, the 50S subunit is in light blue, the mRNA is in blue, and the A- and P-site substrates are colored green and dark blue, respectively. The tripeptidyl moiety of the P- site tRNA is highlighted in yellow. (**B**) 2*Fo-Fc* electron density map (blue mesh) of the ribosome-bound ERY, aminoacyl- and peptidyl-tRNAs in the ribosomal A and P sites, respectively. The refined models of the drug and tRNAs are displayed in their respective electron density maps contoured at 1.0σ. The entire bodies of the A- and P-site tRNAs are viewed from the back of the 50S subunit. Ribosome subunits are omitted for clarity. (**C, D**) Close-up views of the A- and P-site tRNA’s CCA-ends (**C**) and the ribosome-bound ERY (**D**) interacting with nucleotides of the 23S rRNA (blue). H-bonds are shown by black dotted lines. (**E, F**) Close-up views of the tRNAs in the PTC viewed from two different angles, highlighting the characteristic intercalation of the side chain of lysine into the A- site cleft formed by nucleotides A2451 and C2452 of the 23S rRNA (*E. coli* numbering of the rRNA nucleotides is used throughout). Note that -1 Arg of the P-site fMRC-peptidyl-tRNA adopts a conformation that blocks productive accommodation of the lysine residue into the A site.

Except for the apparent differences in the chemical structures of ERY and TEL molecules, the two structures were entirely consistent with each other and nearly indistinguishable. These structures show that the penultimate arginine residue (-1 Arg) of the P-site fMRC-peptidyl-tRNA adopts an unusual conformation in which it partially obstructs the A-site cleft (**Figure 1E, F**), a hydrophobic pocket formed by the 23S rRNA residues A2451, C2452, and U2506 that generally accommodates the side chains of an incoming aa-tRNA. This position is stabilized by the two H-bonds between the side chain of -1 Arg residue and the G2061 and C2452 nucleobases (**Figure 1F**) and is principally similar to one of the two states observed previously for the equivalent arginine residue in the cryo-EM structure of the ERY or TEL-arrested RNCs carrying ErmDL peptide (**Figure S5A, B**)^21^. Repositioning of the -1 Arg residue in the new structures is also confirmed by alignment with the recent structure of the non-arrested RNC containing the same fMRC-peptidyl-tRNA in the P site and Phe-tRNA^Phe^ in the A site (**Figure S6A**)^29^. Importantly, in this conformation, the -1 Arg residue of the nascent peptide would sterically block normal accommodation of the aa-tRNAs with bulky amino acid residues by partially obstructing the A site (**Figure S6A**). Indeed, in both new structures, the lysine moieties of the incoming lysyl-tRNA^Lys^ appear in a previously unseen conformation, in which the lysine side chain is inserted into the A-site cleft but, due to the lack of sufficient space, cannot fully expand and accommodate (**Figure 1C, D; Figure S3C, D**). As a result, the lysine’s α-amino group is positioned at least 4.2 Å away from the carbonyl carbon of the peptidyl-tRNA in the P site, which is too far^34^ for the nucleophilic attack and the peptide bond formation to occur efficiently (**Figure 1C**). Moreover, the observed position of the lysine residue, as well as the A76 ribose pucker of the peptidyl-tRNA, are incompatible with the formation of the proton wire, an intricate network of H-bonds needed for efficient deprotonation of the attacking α-amine during the initial rate-limiting step of peptide bond formation (**Figure S6A**)^35^. At the same time, the CCA-end of the A-site lysyl-tRNA^Lys^ forms canonical interactions with the A-loop (Helix 92) of the 23S rRNA^36^, indicating that the presence of a scrunched lysine residue in the A site does not impede recognition of the CCA-end by the ribosome (**Figure S6A**).

To investigate the proposed allosteric inactivation of the PTC by macrolides, which was suggested to occur through the reorientation of nucleotides A2602 and U2585^20^, we compared the conformations of these key nucleotides in our new macrolide-containing complexes with a previous ribosome structure representing active state of the PTC, carrying the same fMRC-tRNA in the P site but Phe-tRNA instead of Lys-tRNA in the A site^29^. Structural superimposition revealed no major differences in the positions of key functional nucleotides of the 23S rRNA around the PTC, except for a slight displacement of U2585, whose nucleobase was shifted away from the P-site substrate (**Figure S6B**). While this shift might contribute to PTC inhibition, its underlying cause remains ambiguous – whether it results from macrolide binding or simply reflects differences in A-site substrates (Phe-tRNA^Phe^ *vs.* Lys-tRNA^Lys^) remains unclear. Crucially, our data do not support a model in which macrolides allosterically reconfigure the PTC and predispose it to stalling. Instead, the new structures suggest that the inactive conformation of the A-site Lys-tRNA^Lys^ arises primarily due to steric hindrance imposed by the penultimate -1 Arg of the nascent chain rather than a macrolide-induced global rearrangement of the PTC. To definitively address this question, we sought to reconstitute control complexes capturing non-arrested, drug-free RNCs containing the same minimal +X+ motif.

### Macrolides induce stalling-prone conformation of the penultimate arginine

To understand why ribosome stalls at the +X+ motifs only in the presence of a macrolide antibiotic, we determined a 2.6Å-resolution control structure of a non-arrested RNC containing the same fMRC-peptidyl-tRNAi^Met^ and lysyl-tRNA^Lys^ in the P and A sites, respectively, but lacking macrolide bound in the NPET (**Figure 2; Table S1**). It is important to mention that obtaining structures of arrested RNCs does not require the usage of non-hydrolyzable aminoacyl- and peptidyl-tRNAs because PTC is inhibited in such ribosome complexes by default, as exemplified by multiple cryo-EM structures of stalled RNCs, all of which contain native labile ester linkages^15–17,21^. However, obtaining structures of the corresponding non-arrested RNCs has been challenging because the inherent catalytic activity of the PTC leads to rapid transpeptidation and undesired conversion of pre-attack to post-transpeptidation complexes. Thus, to trap non-arrested RNCs in the pre-attack state of the PTC, we used non-hydrolyzable aminoacyl- and peptidyl-tRNAs, in which the labile ester bond is substituted with a stable amide linkage^29^.

**Figure 2.**
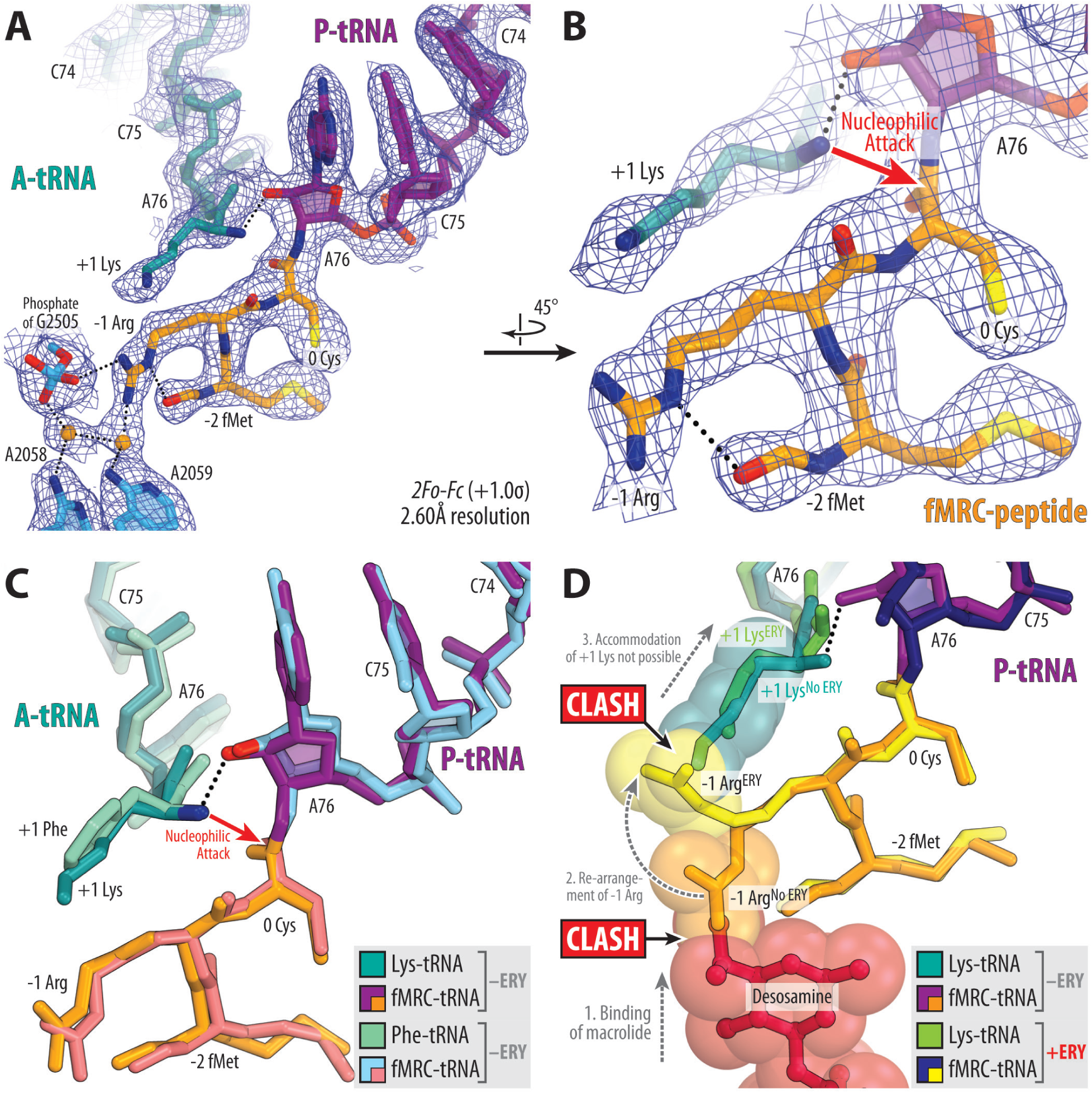
Structure of non-arrested drug-free RNC. (**A, B**) Close-up views of the 2*Fo-Fc* electron density map (blue mesh) of the CCA-ends of Lys-tRNA^Lys^ (teal) and fMRC-peptidyl-tRNAi^Met^ (violet with tripeptidyl moiety highlighted in orange) in the A and P sites, respectively. The refined models of the tRNAs and 23S rRNA nucleotides (blue) are displayed in their respective electron density maps contoured at 1.0σ. H-bonds are shown by black dotted lines. (**C, D**) Superpositions of this new structure of non-arrested macrolide-free RNC with the previous structure of non-arrested 70S ribosome containing A-site Phe-tRNA^Phe^ (greencyan) and the same P-site fMRC-peptidyl-tRNAi^Met^ (light blue with tripeptidyl moiety highlighted in salmon) (**C**, PDB entry 8CVK^29^) or the new structure of arrested RNC containing macrolide ERY (crimson) and the same A- and P-site tRNAs (**D**). Note that, in the absence of NPET-bound macrolide, the side chain of the -1 Arg residue extends away from the A-site cleft and into the NPET, where it interacts with the 23S rRNA. Also, note that desosamine moiety of macrolide clashes with the side chain of -1 Arg, causing its reorientation into the A site, where it interferes with lysine accommodation.

The obtained structure of the drug-free non-arrested RNC shows that the side chain of the -1 Arg residue of the P-site fMRC-peptidyl-tRNA extends away from the A-site cleft and into the NPET, where it establishes electrostatic interactions with the phosphate of G2505 and a water-mediated contact with A2059 (**Figure 2A, B**). This structure featuring Lys-tRNA^Lys^ in the A site is consistent with the previous structure of the non-arrested RNC containing A-site phenylalanyl-tRNA^Phe^ (**Figure 2C**). Moreover, the incoming Lys-tRNA^Lys^ in this structure appears in the canonical pre-attack state, in which its α-amino group is properly positioned relative to the carbonyl carbon of the peptidyl-tRNA for an efficient nucleophilic attack to take place by forming H-bond with the 2’-OH of the peptidyl-tRNA A76 (**Figure 2C**).

Importantly, superpositioning of the structures of non-arrested drug-free and macrolide-arrested RNCs reveals that desosamine moiety of the ribosome-bound ERY (or TEL) sterically overlaps with the extended conformation of the -1 Arg residue of the fMRC-peptidyl-tRNA suggesting that this residue re-orients towards the A site in the presence of a bound macrolide to avoid a clash with it (**Figure 2D; Figure S6C**). The same comparison also revealed no difference in the positions of key PTC nucleobases, suggesting that the previously observed shift of U2585 likely reflects an alteration in response to the binding of different aa-tRNA substrate rather than truly allosteric inactivation of the PTC (**Figure S6B, D**). Therefore, the role of a macrolide in the PTC-stalling mechanism is to induce the specific stalling-prone conformation of the penultimate arginine residue of the nascent peptide, which can interfere with the accommodation of some aa-tRNAs. Furthermore, this new structure demonstrates that the dimethylamino group of a macrolide can directly reach the penultimate position of the 3-amino acid long nascent peptide chain in the NPET (**Figure 2D**), and hence, the rest of the nascent chain might not be necessary for sensing the macrolide. But why must the penultimate residue be a positively charged arginine or lysine for the stalling to occur?

### Penultimate Arg/Lys residues act as macrolide binding sensors in the NPET

It is evident from our structures that, depending on the presence of a macrolide in the NPET, the penultimate arginine acts as a molecular switch toggling between the two alternative conformations: (i) compatible and (ii) incompatible with the incoming aa-tRNAs carrying bulky amino acid residues (such as lysines or arginines). If no macrolide is present, the -1 Arg residue adopts an extended orientation in the NPET, compatible with any incoming tRNAs (**Figure 2C**), and stalling does not occur. Binding of a macrolide to the NPET forces the -1 Arg residue to switch its position to the A site, where it interferes with the productive accommodation of the incoming lysine (and likely arginine) residues (**Figure 2D**), leading to a translational arrest. The key features that allow -1 Arg residue to encroach upon the macrolide binding pocket in the NPET and also to adopt the A-site conformation are its side chain’s length and flexibility: arginines (and lysines) have the longest and the most flexible aliphatic side chains out of all proteinogenic amino acids. The positive charge of the -1 Arg side chain might also contribute to its strong coordination in either of the two conformations. Although we do not have a similar set of structures of arrested and non-arrested RNCs with the lysine residue in the penultimate position of the nascent peptide, *in silico* modeling suggests that the -1 Lys and -1 Arg residues likely behave the same because of their similar length, flexibility, and charge (**Figure 3A**).

**Figure 3.**
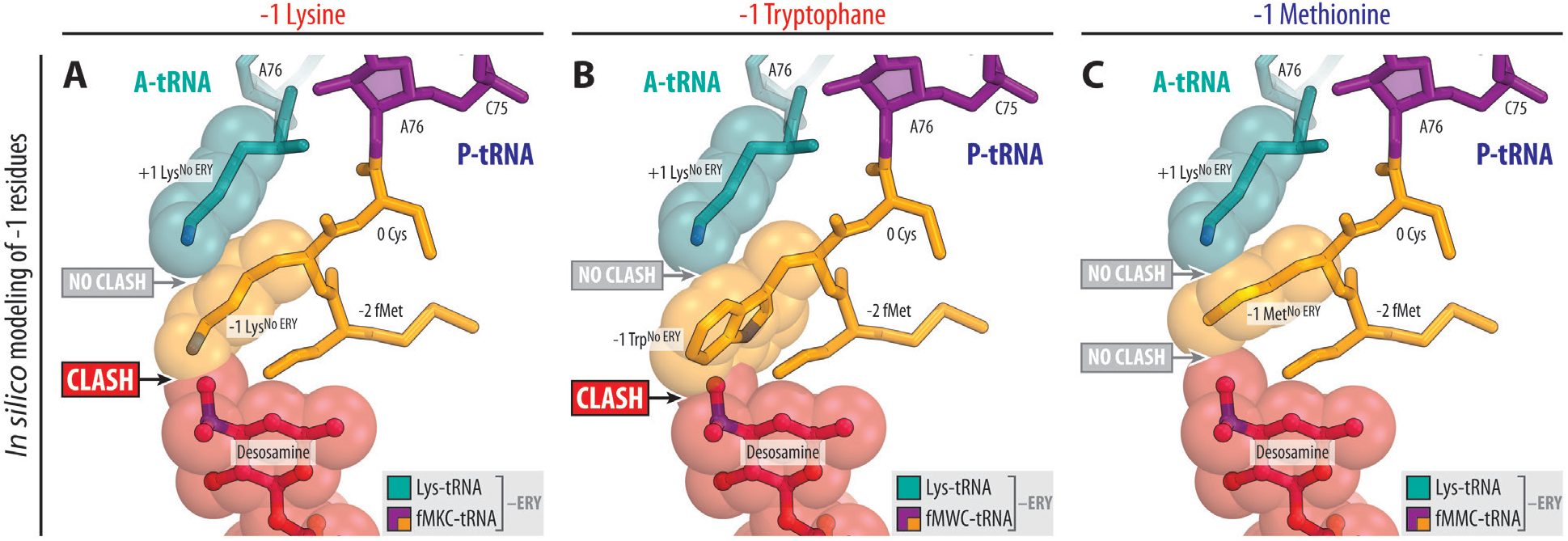
*In silico* modeling of Lys, Trp, or Met residues in the penultimate position of the fMRC-peptidyl-tRNA. Structural models of -1 Lys (**A**), -1 Trp (**B**), and -1 Met (**C**) were generated using the new structure of non-arrested drug-free RNC as a template, with additional reference to other available structures, including the structure with -1 phenylalanine^37^. Note that the desosamine moiety of the macrolide sterically interferes with lysine and bulky aromatic residues (such as tryptophan) in the penultimate position of the nascent chain but not with smaller side chains like methionine.

Why, then, do other bulky residues in this position, such as tryptophane, tyrosine, or phenylalanine, fail to induce ribosome stalling? *In silico* modeling based on the known structure of phenylalanine in the penultimate position^37^ suggests that bulky aromatic residues, due to their limited flexibility and the spatial constraints of the PTC around the nascent peptide, cannot bend (like Arg or Lys can) and adopt a conformation that obstructs the A site. Instead, these residues likely adopt only a single rotameric conformation resembling the extended NPET orientation of the -1 arginine residue (**Figure 3B**). In this orientation, the -1 Trp/Tyr/Phe side chain would not interfere with the aa-tRNA accommodation into the A site, explaining why stalling does not occur. However, these residues sterically clash with the desosamine moiety of the macrolide (**Figure 3B**), likely displacing the drug. As a result, bulky aromatic side chains in the penultimate position do not cause ribosome stalling because they cannot re-orient into the A-site cleft like -1 Arg or -1 Lys residues can. Conversely, smaller side chains of other possible -1 residues, regardless of their flexibility, are too short to interfere with macrolide binding in the NPET (**Figure 3C**).

To test this hypothesis experimentally, we determined a structure of non-arrested RNC containing NPET-bound ERY, A-site lysyl-tRNA^Lys^, and P-site fMAC-peptidyl-tRNAi^Met^, in which penultimate arginine residue is replaced with alanine (**Figure 4A; Table S1**). The PTC in this structure appears in the canonical pre-attack state with a fully accommodated lysine residue into the A site (**Figure 4B, C**), consistent with the observed lack of ribosome stalling on the MACK sequence in the toe-printing assay (**Figure S2**, lane 8). Superpositioning of this structure with the corresponding arrested RNC revealed no substantial differences in the overall conformations of the fMet-Ala-Cys *vs.* fMet-Arg-Cys peptide chains of the peptidyl-tRNAs in the NPET (**Figure 4D**). Moreover, not only do the trajectories of these tripeptides look the same, but even the positions of the main-chain and Cβ atoms of -1 Ala or -1 Arg are identical (**Figure 4D**). Altogether, these data suggest that the unique combination of flexibility and size endow the penultimate Arg/Lys residue of the nascent peptide chain with the ability to act as a macrolide binding sensor that transmits the signal from the NPET to the A site.

**Figure 4.**
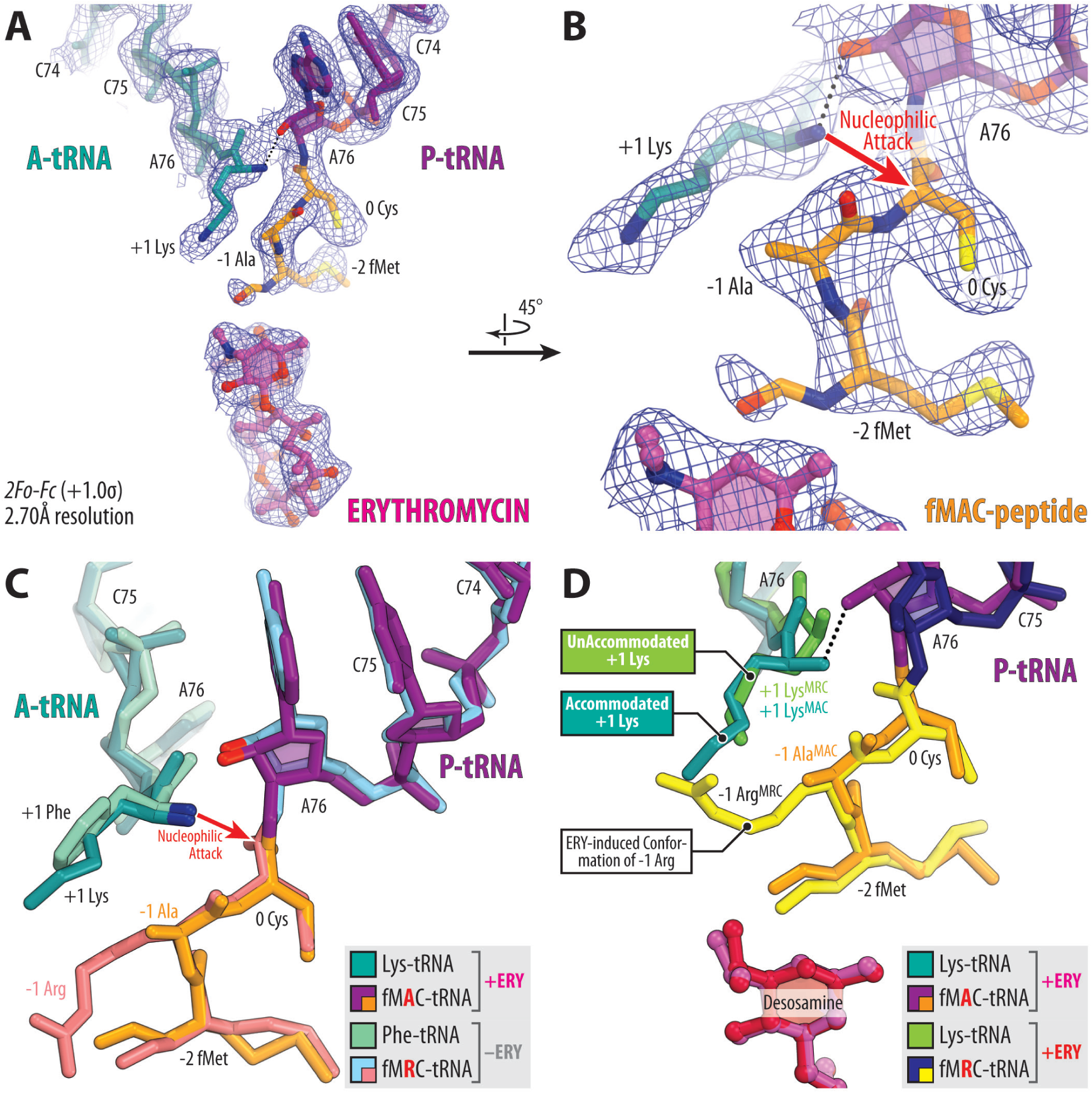
Structure of non-arrested RNC with penultimate arginine mutated to alanine. (**A, B**) Close-up views of the 2*Fo-Fc* electron density map (blue mesh) of the ribosome-bound ERY (magenta) and the CCA-ends of Lys-tRNA^Lys^ (teal) and mutated fMAC-peptidyl-tRNAi^Met^ (violet with tripeptidyl moiety highlighted in orange) in the A and P sites, respectively. The refined models of the drug and tRNAs are displayed within their respective electron density maps contoured at 1.0σ. (**C, D**) Structural alignments comparing this new non-arrested macrolide-bound RNC with previously reported structures. (**C**) Superimposition with the non-arrested 70S ribosome containing A-site Phe-tRNA^Phe^ (greencyan) and P-site fMRC-peptidyl-tRNAiMet (light blue, with the tripeptidyl moiety in salmon) (PDB entry 8CVK^29^). (**D**) Comparison with the new structure of arrested RNC containing ERY (crimson). Note that the absence of the -1 Arg residue removes steric hindrance, allowing both macrolide binding and proper accommodation of lysyl-tRNA into the PTC.

### Incoming non-Lys/Arg residues prevent ribosome stalling via distinct mechanisms

Our structural analysis shows that macrolide-induced conformation of the -1 Arg/Lys residue in the growing polypeptide renders A site of the PTC selective against the incoming +1 Arg/Lys residue, resulting in translational arrest. However, it is unclear why A site rejects only the incoming lysine or arginine but not other residues^38^. *In silico* modeling and molecular dynamics simulations^21^ suggested that macrolide-induced conformation of -1 Arg should interfere not only with the accommodation of the incoming +1 Arg/Lys residue but also with the equally large aromatic side chains, such as Phe, Tyr, or Trp. Nevertheless, substituting the +1 lysine residue in the +X+ motif sequence with phenylalanine (**Figure S2**, lane 10) or other aromatic residues^38^ completely abolishes macrolide-dependent ribosome stalling, indicating that these residues accommodate into the A site, overcome the occlusion imposed by the -1 Arg, and react with the P-site peptidyl-tRNA just normally. The same effect is also observed if the +1 residue is changed to a proline (**Figure S2**, lane 12) or other relatively small residues^38^ that are not expected to clash with the macrolide-induced conformation of -1 Lys/Arg residue.

To understand why only +1 Lys/Arg, among all bulky amino acid residues, induce ribosome stalling at the +X+ motifs, we determined a structure of non-arrested RNC containing the fMRC-peptidyl-tRNAi^Met^ in the P site and aminoacylated phenylalanyl-tRNA^Phe^ in the A site at 2.40Å resolution (**Figure 5A, B**). Consistent with other structures of non-arrested RNCs reported here, the A- and P-site substrates in these complexes appear in the canonical pre-attack state, with aromatic Phe residue clearly resolved in the electron density map and fully accommodated into the A-site cleft (**Figure 5B**). Surprisingly, although ERY was also included in the crystallization mixture, no electron density for the ribosome-bound macrolide was detected within the NPET. Additional experiments using other macrolides, including TEL or AZI, and higher drug concentrations yielded the same result – no visible density for the macrolide (**Figure 5A**). This suggests that the accommodation of bulky residues, such as Phe, into the A site displaces macrolides from the tunnel. Because these structures lack ribosome-bound macrolides and are nearly indistinguishable from the recently published structure of non-arrested RNC (PDB entry 8CVK)^29^, the corresponding data were not deposited in the PDB.

**Figure 5.**
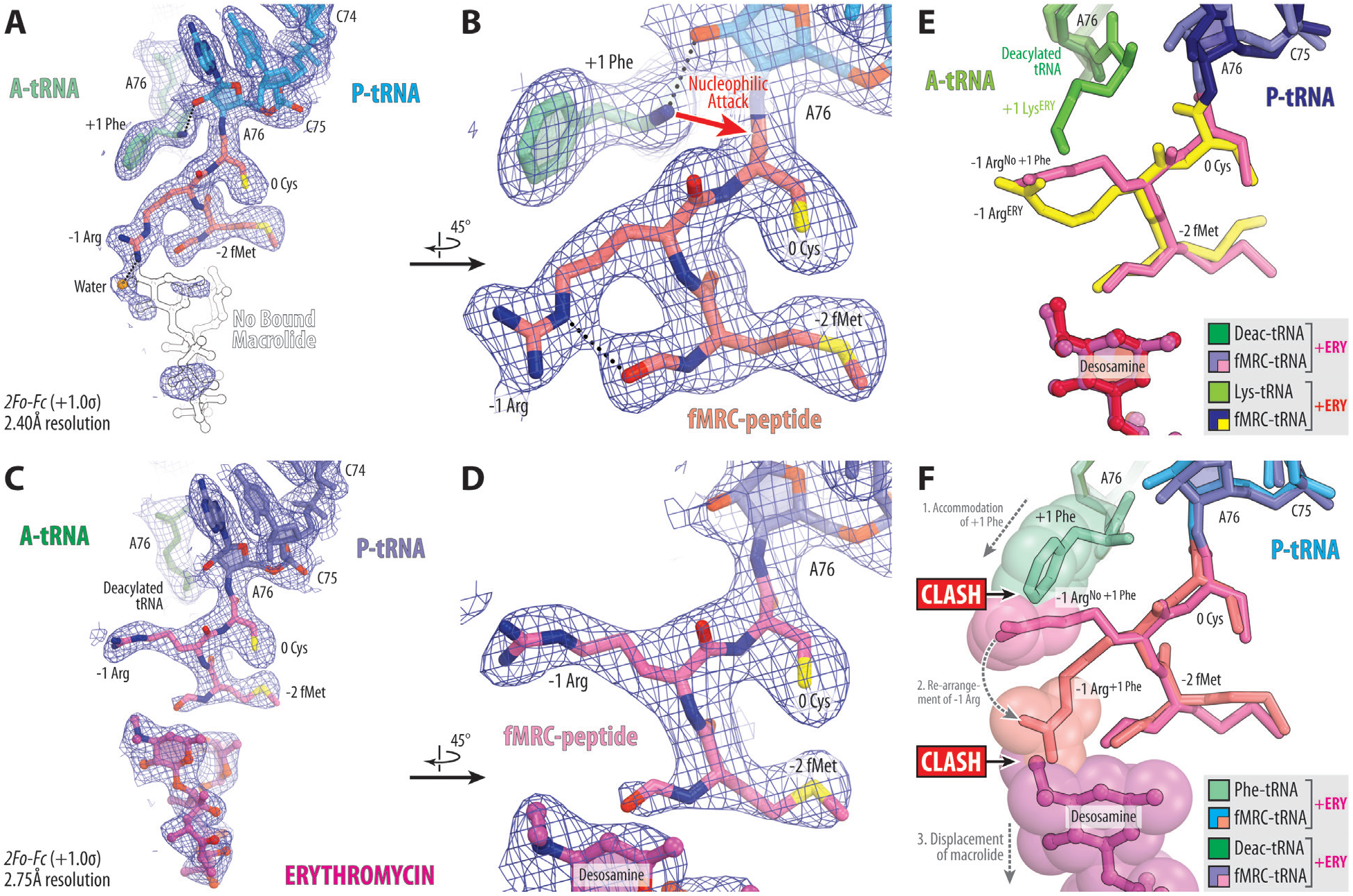
Structural analysis of non-arrested RNCs with modified A-site substrates. (**A-D**) Close-up views of the 2*Fo-Fc* electron density map (blue mesh) of the macrolide binding site and the CCA-ends of the P-site fMRC-peptidyl-tRNAi^Met^ (blue/slate with tripeptidyl moiety highlighted in salmon/pink) and either aminoacylated Phe-tRNA^Phe^ (**A, B**, greencyan) or deacylated tRNA^Phe^ (**C, D**, dark green) in the A site. In panel **A**, the lack of macrolide binding is highlighted by a faint contour. In panels **C** and **D**, ribosome-bound ERY is shown in magenta. The refined models of the drug and tRNAs are displayed within their respective electron density maps contoured at 1.0σ. (**E**) Superimposition of the structures of the new non-arrested macrolide-bound RNC with deacylated tRNA in the A- site with arrested RNC containing ERY (crimson). Note that macrolide binding in the NPET repositions the -1 Arg residue of the nascent chain towards the A site. (**F**) Structural comparison between two non-arrested RNCs with either aminoacylated or deacylated tRNA^Phe^ in the A site. Note that the accommodation of +1 Phe and the subsequent rearrangement of -1 Arg lead to a steric clash with the macrolide, resulting in its displacement from the ribosomal tunnel.

Similar to other structures of non-arrested RNCs, the -1 Arg residue of the fMRC-peptidyl-tRNA in this structure appears in the extended conformation incompatible with macrolide binding and likely acts as a “middleman,” transmitting the signal in the reverse direction, from the A site to the macrolide-binding pocket in the NPET. To establish a causal link between the presence of the bulky aromatic +1 Phe residue in the A site and the absence of macrolide in the NPET, we determined the structure of a similar 70S ribosome complex containing A-site deacylated tRNA^Phe^ at 2.75Å resolution (**Figure 5C, D; Table S1**). In this structure, the ribosome-bound ERY was clearly resolved in the NPET, and the -1 Arg of the peptidyl-tRNA adopted the macrolide-induced A-site conformation, not entirely identical but similar to the one observed in the ERY-stalled RNC (**Figure 5E**). While deacylated tRNA^Phe^ is not a physiologically relevant A-site substrate, here we used it merely as a mimic for tRNA with smaller amino acid residues, contrasting with bulky Lys/Arg or aromatic residues. Although any incoming non-Lys/Arg residue prevents ribosome stalling on +X+ motifs^38^, our structural data suggest two distinct mechanisms by which different aa-tRNAs counteract the inhibitory effects of macrolide antibiotics: (i) amino acids with small side chains avoid overlap with the -1 Lys/Arg of the peptidyl-tRNA and can be properly accommodated into the ribosomal A site even in the presence of NPET-bound macrolides; and (ii) bulky aromatic residues, while also normally accommodated in the A site, clash with the -1 Lys/Arg of the peptidyl-tRNA, physically displacing macrolides from the ribosome (**Figure 5F**).

### The mechanism of stalling at the +X+ motif (DISCUSSION)

The detailed structural analysis of arrested and non-arrested RNCs presented here uncovers the roles of each individual component in macrolide-induced ribosome stalling at the minimal +X+ motif. We show that binding of a macrolide in the NPET induces a stalling-prone conformation of the penultimate Arg or Lys residue in the growing polypeptide chain. These residues act as sensors for macrolide binding, transmitting a signal to the ribosomal A site through their long, flexible side chains. Macrolide-induced change in the conformation of -1 Arg/Lys residues alters the selectivity of the A site, which can no longer accommodate incoming +1 Arg/Lys residues, leading to ribosome stalling and translational arrest (**Figure 6A**).

**Figure 6.**
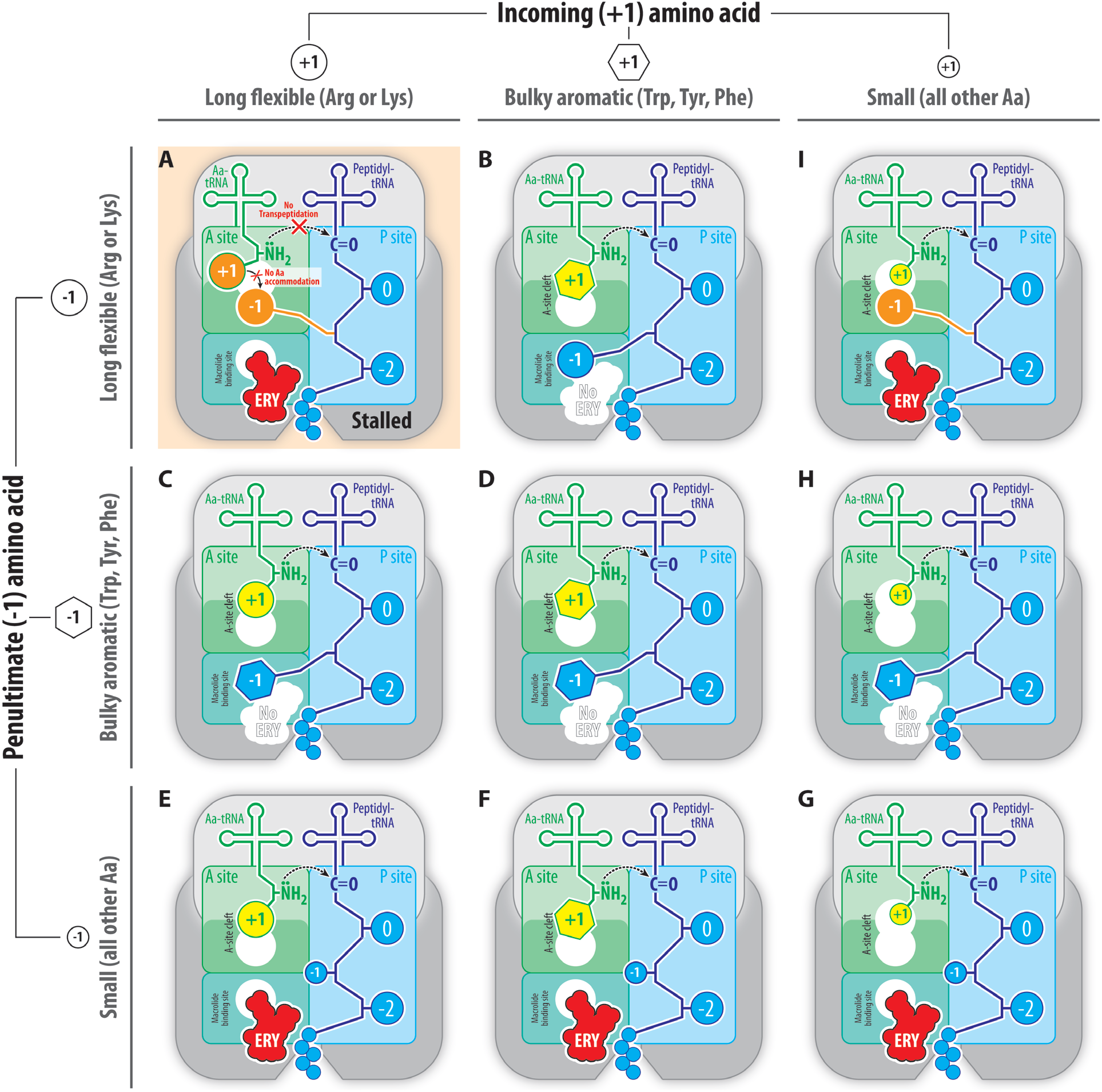
Mechanism of macrolide-induced ribosome stalling at the +X+ motifs. (**A-I**) Schematic diagram illustrating the outcomes of different combinations of -1 and +1 amino acid residues present in the PTC together with macrolide antibiotics. The macrolide (red), the penultimate (-1) residue of the nascent peptide chain (orange), and the incoming (+1) amino acid (blue) are highlighted. (**A**) Stalling conditions. Ribosome stalling on +X+ motifs occurs when macrolide-induced conformation of the -1 Arg/Lys residue of the growing polypeptide chain sterically hinders productive accommodation of the +1 Arg/Lys residue into the A site, making peptide bond formation impossible. (**B-I**) Non-stalling conditions. Several alternative -1 and +1 residue combinations allow translation to proceed: (**B**) when the -1 residue of the nascent peptide is Arg/Lys and the +1 residue contains a bulky aromatic side chain or (**C, D, H**) if the -1 residue is bulky aromatic, translation continues regardless of the identity of the +1 residue, as macrolides are displaced, and aminoacyl-tRNA successfully accommodates into the A site; (**E, F, G**) when the -1 residue lacks a large side chain (i.e., residues other than Arg/Lys/Trp/Tyr/Phe/His), there is no steric clash with the macrolide or the A-site tRNA, allowing normal peptide bond formation; (**I**) similarly, if the +1 residue is small, steric hindrance is minimized, and translation proceeds unimpeded. This diagram shows how the interplay between ribosome-bound macrolide (red), the -1 (orange), and +1 (yellow) residues determine ribosome stalling at the +X+ motifs.

In the other possible combinations of +X+ stalling determinants, aa-tRNAs normally accommodate into the PTC and react with the peptidyl-tRNA so that stalling does not occur (**Figure 6B-I**). However, depending on the identity of the penultimate (-1) or the incoming (+1) residues, the actual mechanism allowing the ribosome to bypass the stalling could be different. When the -1 residue of the nascent peptide is Arg/Lys and the +1 residue contains a bulky aromatic side chain (**Figure 6B**), or if the -1 residue is bulky aromatic regardless of the identity of the +1 residue, translation continues because macrolides are displaced, and aa-tRNAs can productively accommodate into the A site and react with a peptidyl-tRNA on the drug-free ribosome (**Figure 6C, D, H**). When the - 1 residue lacks a large side chain (i.e., residues other than Arg/Lys/Trp/Tyr/Phe/His), there is no steric clash with the macrolide or the A-site tRNA, allowing normal peptide bond formation (**Figure 6E, F, G**). Similarly, if the +1 residue is small, steric hindrance is minimized, and translation proceeds unimpeded (**Figure 6I**). Importantly, in this work, we show that the macrolide desosamine group – and, more specifically, its dimethyl-amino moiety – is crucial not only for drug binding to the ribosome^33,39^ but also for mediating the macrolide-induced repositioning of the -1 Arg/Lys residues in the peptidyl-tRNA. Since all clinically used macrolides contain the same dimethyl-amino group, the mechanism of ribosome stalling at +X+ motifs revealed here is likely to be conserved across most macrolides, explaining why stalling persists despite alterations to the drug’s chemical structure outside of this moiety.

However, a lingering question is why +1 Lys/Arg residues cannot push the -1 Lys/Arg residue of the peptidyl-tRNA to displace the macrolide, as Phe does (**Figure 5F**). We hypothesize that this difference arises from the varying affinities of aromatic versus aliphatic residues (Lys/Arg) for the A-site cleft. Bulky aromatic amino acids (e.g., Phe, Tyr, Trp, or His) snugly fit into the hydrophobic pocket formed by nucleobases A2451 and C2452, establishing strong π-π stacking interactions and, thus, high binding affinities. In contrast, aliphatic Lys/Arg residues can only form weaker CH-π interactions with these nucleobases. While these affinity differences may not significantly affect normal translation, the presence of a macrolide in the NPET, combined with the reorientation of -1 Lys/Arg residue in the growing polypeptide, creates an insurmountable energetic barrier for accommodation of +1 Arg/Lys, rendering them unreactive. The outcome ultimately depends on which entity – the macrolide or the aa-tRNA – has the stronger affinity for the ribosome, resulting in two possible scenarios: (i) the macrolide remains bound in the NPET while aa-tRNA cannot be accommodated (**Figure 6A**), or (ii) the bulky aromatic side chain of -1 residue displaces the macrolide (**Figure 6C, D**). Importantly, this competition between the NPET-bound macrolide and the incoming aa-tRNA is indirect, occurring through the -1 Lys/Arg residue of the nascent polypeptide chain. Therefore, if the -1 Lys/Arg residue is absent, no stalling occurs, as aa-tRNA can fully accommodate into the A site regardless of the side chain size (**Figure 6E, F, G**). Similarly, any +1 residues other than Arg/Lys or aromatic Phe/Tyr/Trp/His can readily accommodate into the A site and react with the peptidyl-tRNA, regardless of macrolide presence or the identity of the -1 residue (**Figure 6G, H, I**).

This study focuses on the shortest and the most robust MAM sequence, MRCK, representing the +X+ consensus in the purest form, enabling a detailed analysis of each stalling determinant. In cellular contexts, however, most +X+ stalling sites occur within longer nascent chains, where upstream sequences can modulate stalling efficiency^11,18^. The effects range from MAM inactivation^11^ to changes in the stalling mechanism depending on the macrolide^21^. For instance, the +X+ motif in the ErmD leader peptide (MTHSM**R**L**R**) retains its stalling ability in response to TEL only while its +X+ identity is preserved^21^. In contrast, ERY stalls translation at ErmDL even when both -1 and +1 arginines are mutated to alanines^21^, suggesting that stalling determinants in the full-length ErmDL peptide are present not only in its C-terminal +X+ motif but also in its N-terminal region. Likely, this ERY-induced, (+X+)-independent stalling arises from interactions between ERY’s cladinose sugar and the N-terminal portion of the ErmDL peptide, which promotes the inactive state of the PTC – a mechanism absent with TEL lacking cladinose.

Leader peptides regulating the expression of macrolide-resistance methyltransferases, such as ErmDL, have evolved to sense low macrolide concentrations, representing quite unique and efficient stalling sequence. However, most +X+-containing motifs identified in ribo-seq data lack ErmDL-like N-terminal consensus sequence, suggesting that cladinose-containing macrolides, such as ERY or AZI, primarily induce stalling through a +X+-mediated mechanism, as described here. Moreover, cladinose-lacking ketolides, such as TEL, rely solely on direct (+X+)-mediated stalling. Indeed, analysis of the ribo-seq data confirms that both ERY and TEL stall ribosomes at the same +X+ sites (**Figure S7**, red), while ERY- or TEL-specific arrest sites are less common (**Figure S7**, green and blue), supporting the generalizability of the described stalling mechanism.

Since the discovery of selective protein synthesis inhibition^11^, the concept of modulating the selectivity of existing inhibitors – or designing novel molecules to target specific proteins – has gained significant attention. This approach holds particular promise for inhibitors of eukaryotic translation, where selectively suppressing harmful protein synthesis could open new avenues for treating human diseases. While some progress has been made^40,41^, the limited understanding of the context-specificity of current inhibitors in both bacteria and eukaryotes presents a significant barrier to further advancements. Importantly, recent studies on macrolide-sensitized *Saccharomyces cerevisiae* have illuminated +X+ as the main drug-arrest motif in eukaryotes as well^22^. We believe that further investigation of a more subtle, broader context that affects the strength of drug-induced stalling at the +X+ motifs will guide knowledge-based development of principally new medicines possessing protein-specific action. Additionally, our incomplete knowledge of how nascent chains fold co-translationally within the NPET complicates the interpretation of structural and biochemical data on selective ribosome inhibition. Encouragingly, the growing body of research into selective ribosome inhibition continues to uncover new molecular insights, advancing our understanding of translational stalling mechanisms.

## ACKNOWLEDGMENTS

We thank Drs. Nora Vázquez-Laslop, Alexander Mankin, and Dmitrii Travin for critical reading of the manuscript and insightful suggestions. We thank the staff at NE-CAT beamlines 24ID-C and 24ID-E for help with X-ray diffraction data collection, especially Drs. Malcolm Capel, Frank Murphy, Surajit Banerjee, Igor Kourinov, David Neau, Jonathan Schuermann, Narayanasami Sukumar, Anthony Lynch, James Withrow, Kay Perry, Ali Kaya, and Cyndi Salbego.

This work is based upon research conducted at the Northeastern Collaborative Access Team beamlines (24ID-C and 24ID-E), which are funded by the National Institute of General Medical Sciences from the National Institutes of Health [grant P30-GM124165 to NE-CAT]. The Eiger 16M detector on 24ID-E beamline is funded by an NIH-ORIP HEI grant [S10-OD021527 to NE-CAT]. This research used resources of the Advanced Photon Source, a US Department of Energy (DOE) Office of Science User Facility operated for the DOE Office of Science by Argonne National Laboratory under Contract No. DE-AC02-06CH11357.

This work was supported by the National Institute of General Medical Sciences of the National Institutes of Health [grants R01-GM132302 and R35-GM151957 to Y.S.P.], Illinois State startup funds [to Y.S.P.], and National Science Foundation [MCB-2345351 to M.S.S.]. The funders had no role in study design, data collection and analysis, decision to publish, or manuscript preparation.

## AUTHOR CONTRIBUTIONS STATEMENT

E.A.S. and E.V.A. prepared non-hydrolyzable full-length peptidyl-tRNAs; E.V.A., A.A.K., and M.N.P. prepared non-hydrolyzable aminoacyl-tRNAs; E.A.S. and M.S.S. performed *in vitro* toe-printing assays; E.A.S., E.V.A., A.A.K., M.N.P., and Y.S.P. designed and performed structural-biology experiments; M.S.S. and Y.S.P. supervised the experiments. All authors interpreted the results. E.A.S., M.S.S., and Y.S.P. wrote the manuscript.

## SUPPLEMENTARY FIGURES

**Figure S1.**
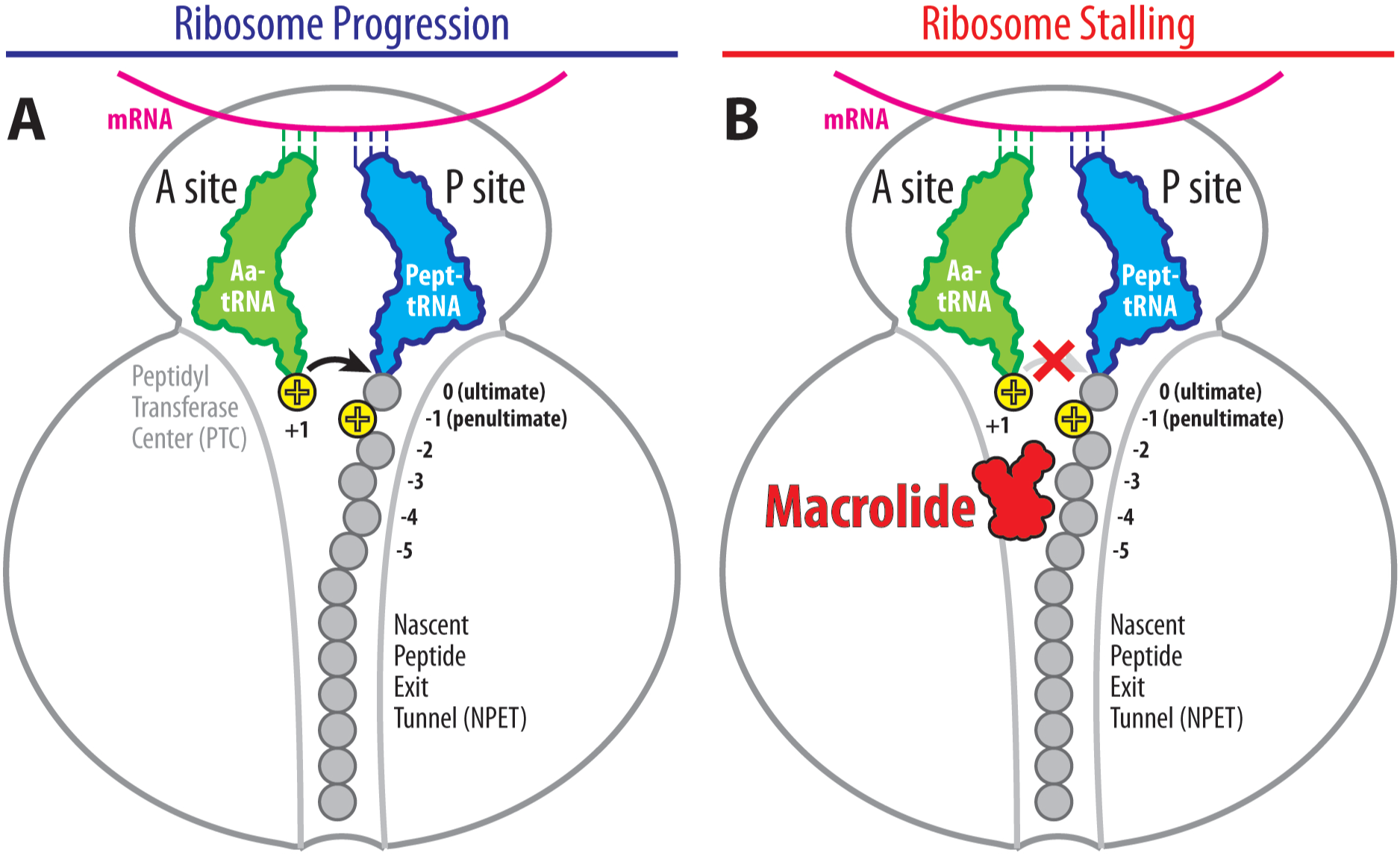
Context-specificity of macrolide action. (**A**) Normal progression of the translating ribosome through +X+ motifs (or any other sequences) in the absence of a macrolide. Peptide bond formation in the PTC occurs normally (black arrow). (**B**) Macrolide-induced inhibition of peptide bond formation in the PTC of a ribosome progressing through +X+ sequence (red cross). Note that NPET-bound macrolide (red) is too far to establish direct contact with the incoming Arg/Lys residues (yellow) in the A site.

**Figure S2.**
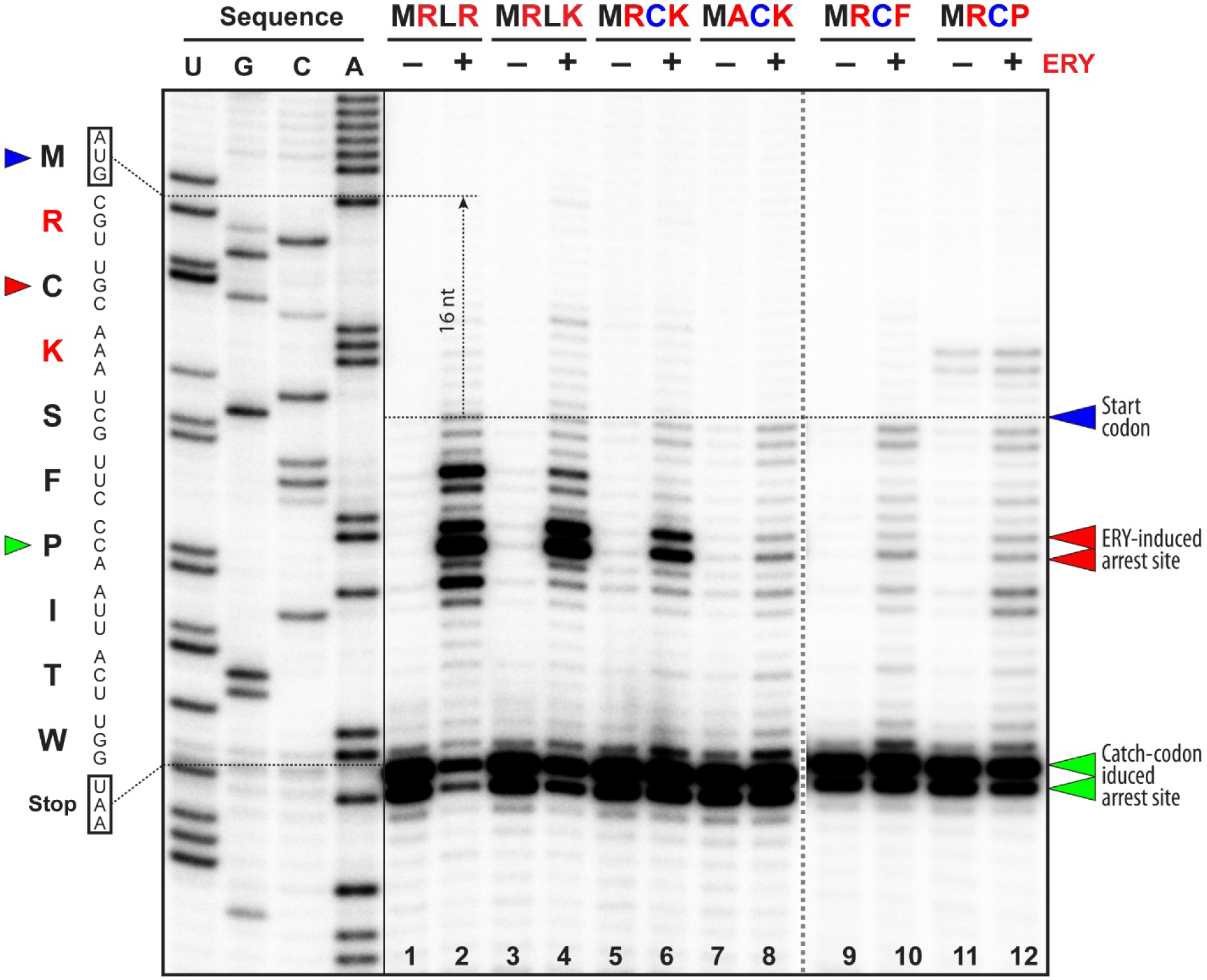
Macrolide ERY arrests translation on MRCK peptide sequence. Ribosome stalling in the presence and absence of ERY revealed by reverse-transcription primer-extension inhibition (toe-printing) assay in a cell-free translation system on mRNA encoding MRLR peptide sequence at the N-terminus (lanes 1-2) or its mutant versions encoding varios amino acids in +1, 0 and -1 positions (lanes 3-12). Nucleotide sequence of MRCK mRNA and the corresponding amino acid sequence are shown on the left. Blue arrowhead marks translation start site. Red arrowheads point to the ERY-induced arrest site within the coding sequences of each of the used mRNAs. Note that due to the large size of the ribosome, the reverse transcriptase used in the toe-printing assay stops 16 nucleotides downstream of the codon located in the P-site. The toe-printing reactions were also supplemented with mupirocin, an inhibitor of isoleucyl-tRNA-synthetase, to arrest all translating ribosomes that bypassed drug-specific stall sites at the downstream isoleucyl codon (green arrowheads). Note that the presence of MRL- or MRC-peptidyl-tRNAs in the P site and arginyl- or lysyl-tRNAs in the A site sustains ERY-dependent stalling, whereas MAC-tripeptide sequence or phenylalanine or proline as incoming residues are not conducive to ribosome stalling and serve as negative controls. Experiments were repeated twice independently with similar results.

**Figure S3.**
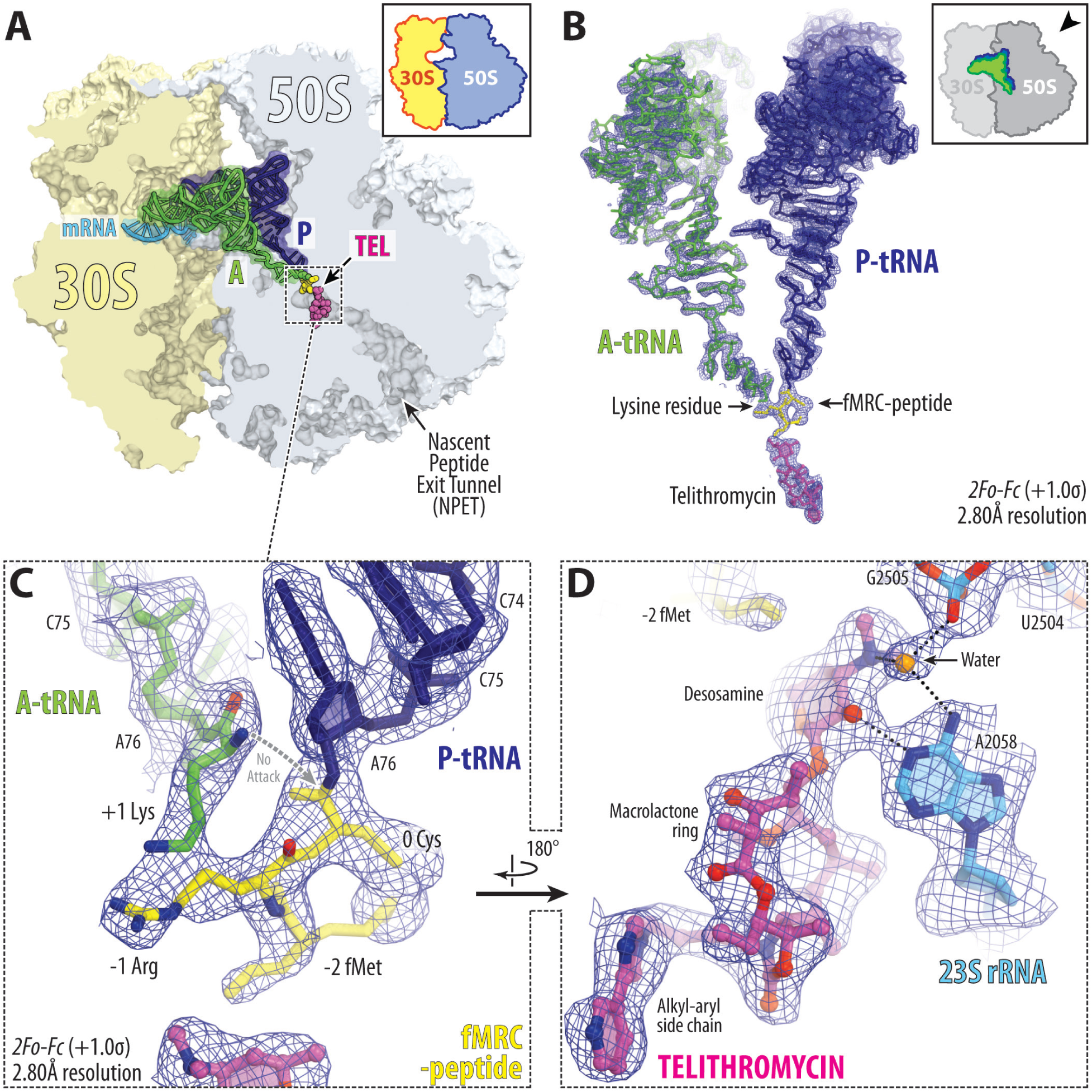
Structure of the telithromycin-arrested RNC assembled from purified components. (**A**) Overview of the structure of the *T. thermophilus* 70S ribosome in complex with telithromycin (TEL, magenta) and also containing non-hydrolyzable lysyl-tRNA^Lys^ and fMRC-peptidyl-tRNAi^Met^ in the A and P sites, respectively. The 30S subunit is shown in light yellow, the 50S subunit is in light blue, the mRNA is in blue, and the A- and P-site substrates are colored green and dark blue, respectively. The fMRC-tripeptidyl moiety of the P-site tRNA is highlighted in yellow. (**B-D**) 2*Fo-Fc* electron density map (blue mesh) of the ribosome-bound TEL, aminoacyl- and peptidyl-tRNAs in the ribosomal A and P sites, respectively. The refined models of the drug and tRNAs are displayed in their respective electron density maps contoured at 1.0σ. The entire bodies of the A- and P-site tRNAs are viewed from the back of the 50S subunit. Ribosome subunits are omitted for clarity. (**C, D**) Close-up views of the A- and P-site tRNA’s CCA-ends (**C**) and the ribosome-bound TEL (**D**). H-bonds are shown by black dotted lines.

**Figure S4.**
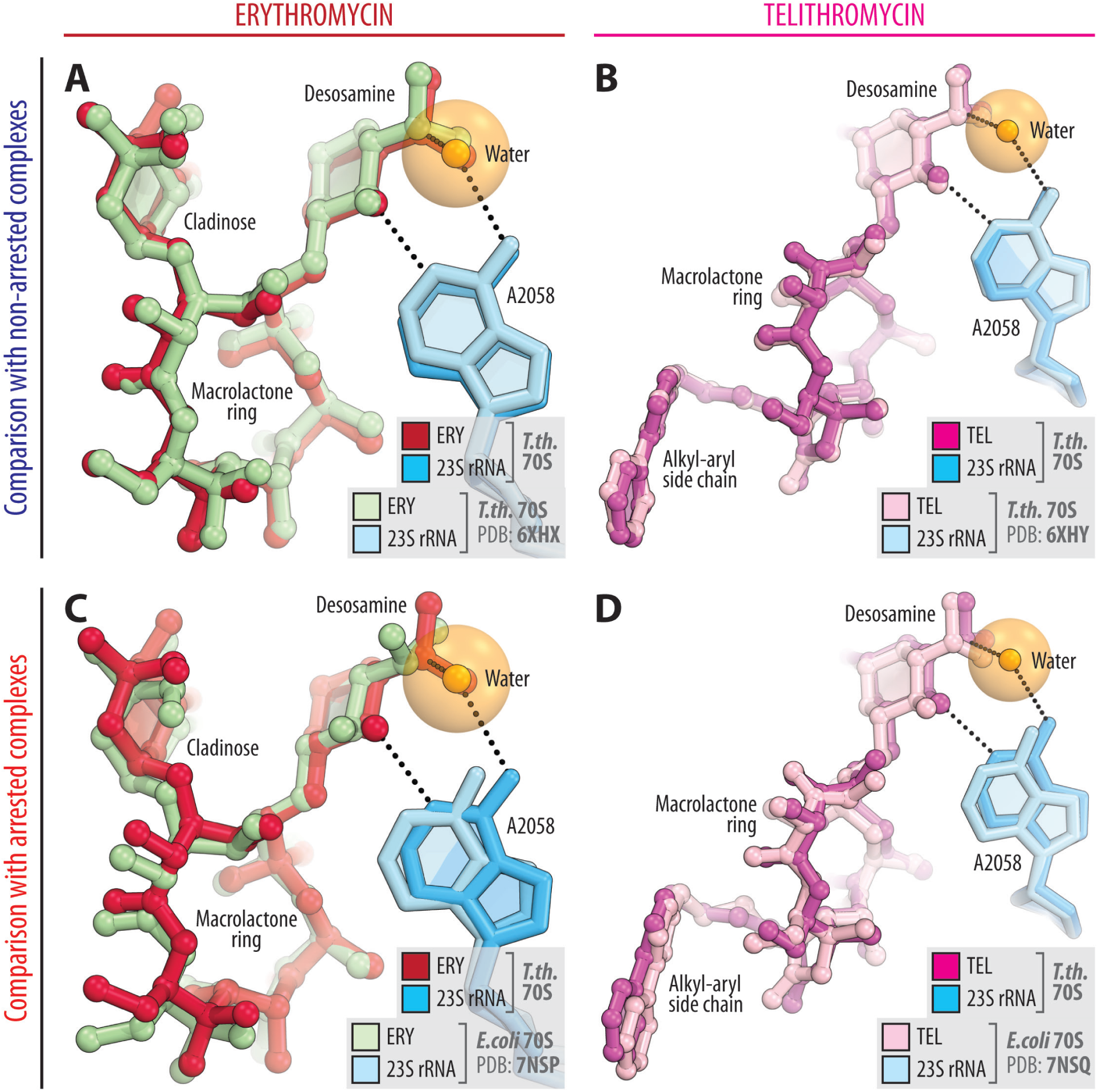
Comparison of macrolide binding sites in the new and previous structures. (**A, B**) Superpositions of the new structures of arrested RNCs containing erythromycin (A, C, crimson) or telithromycin (**B, D**, magenta) with the previous structures of the same drugs (greencyan and pink, respectively) observed in the context of non-arrested *Tth* 70S ribosome (**A, B**, PDB entries 6XHX or 6XHY^33^) or arrested *E. coli* 70S RNCs (C, D, PDB entries 7NSP or 7NSQ^21^). All structures were aligned based on domain V of the 23S rRNA. Note that the binding sites of ERY and TEL are nearly identical regardless of whether the drugs are bound to the arrested or non-arrested RNCs.

**Figure S5.**
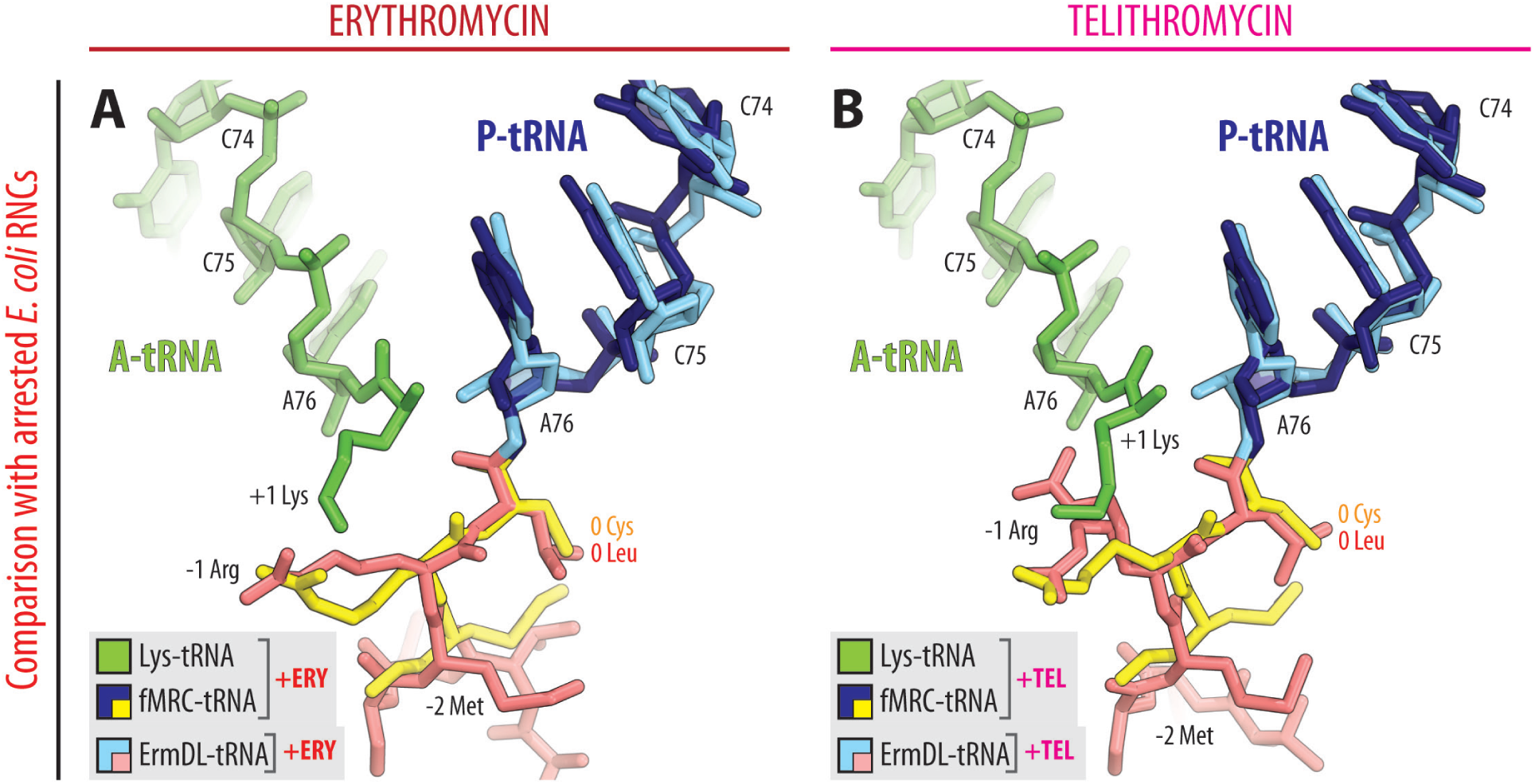
Comparisons of the peptidyl-tRNA conformations in the new and previous stalled RNC structures. (**A, B**) Superpositions of the new structures of ERY or TEL-arrested RNCs carrying fMRC-peptidyl-tRNAs in the P site (dark blue with tripeptidyl moiety highlighted in yellow) with the previous structures of similar *E. coli* 70S RNCs carrying full-length MTHSMRC-peptidyl-tRNAs in the P site (light blue with tripeptidyl moiety highlighted in salmon). All structures were aligned based on domain V of the 23S rRNA. Note that the unusual conformations of the -1 Arg side chain in the short MRC peptides are principally similar to those observed previously for the equivalent -1 Arg residue in the RNCs carrying ErmDL peptides (PDB entries 7NSP or 7NSQ^21^).

**Figure S6.**
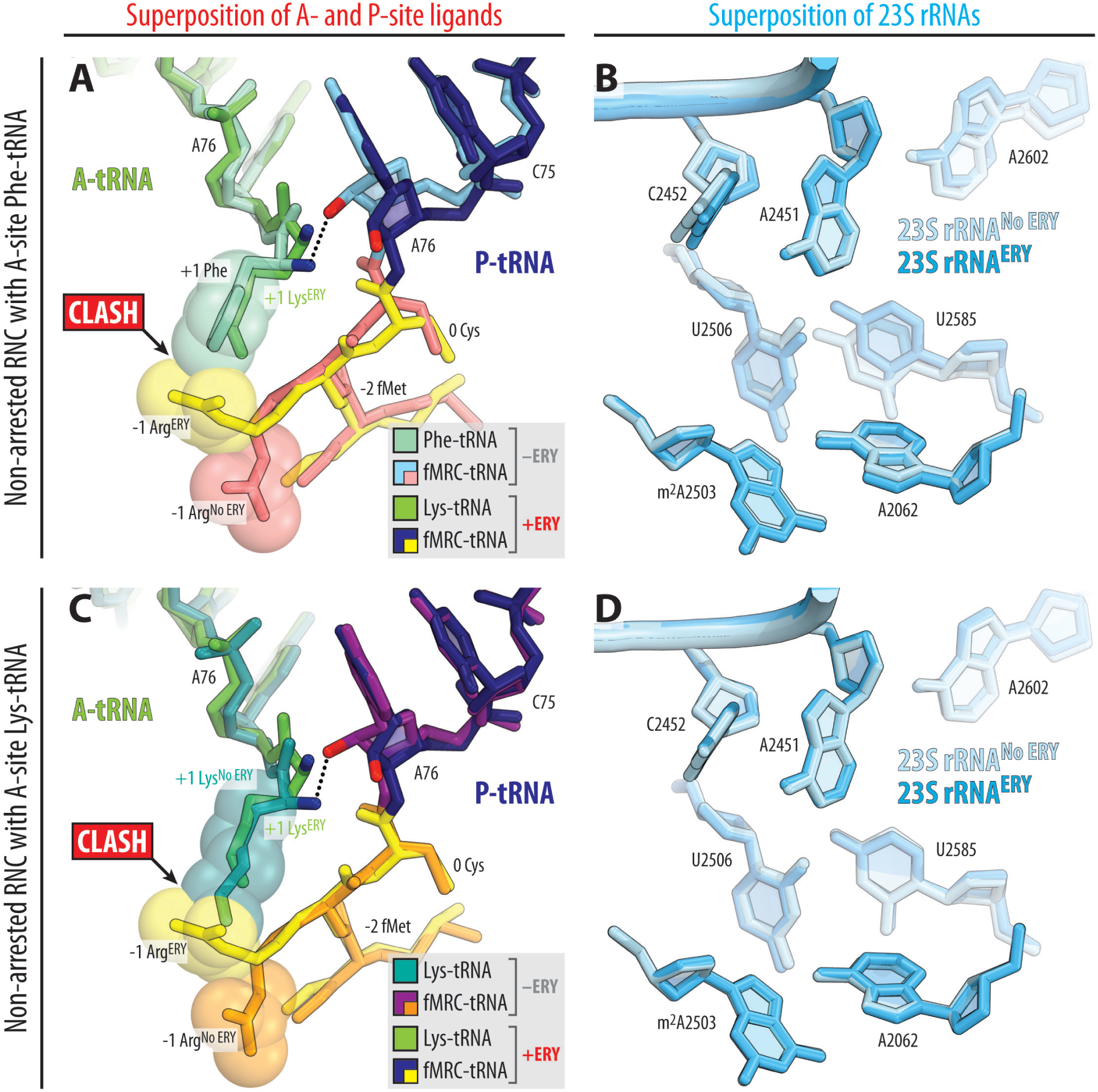
Comparison of the structures of macrolide-arrested RNC and non-arrested RNCs. Superpositioning of the new structure of ERY-arrested RNC carrying Lys-tRNA^Lys^ (green) and fMRC-peptidyl-tRNAi^Met^ (dark blue with tripeptidyl moiety highlighted in yellow) in the A and P sites, respectively, with the previous structure of non-arrested (lacking macrolide) *Tth* 70S ribosome containing A-site Phe-tRNA^Phe^ (greencyan) and the same P-site fMRC-peptidyl-tRNAi^Met^ (light blue with tripeptidyl moiety highlighted in salmon) (**A, B**; PDB entry 8CVK^29^) or the new structure of non-arrested RNC containing the same A- and P-site tRNAs (**C, D**). All structures were aligned based on domain V of the 23S rRNA. (**A, C**) Comparisons of the positions of A- and P-site substrates in the PTC. (**B, D**) Comparisons of the positions of key 23S rRNA nucleotides around the PTC. Note that, in the ERY-arrested RNC, the -1 Arg residue adopts a conformation that interferes with the accommodation of aa-tRNAs into the A site.

**Figure S7.**
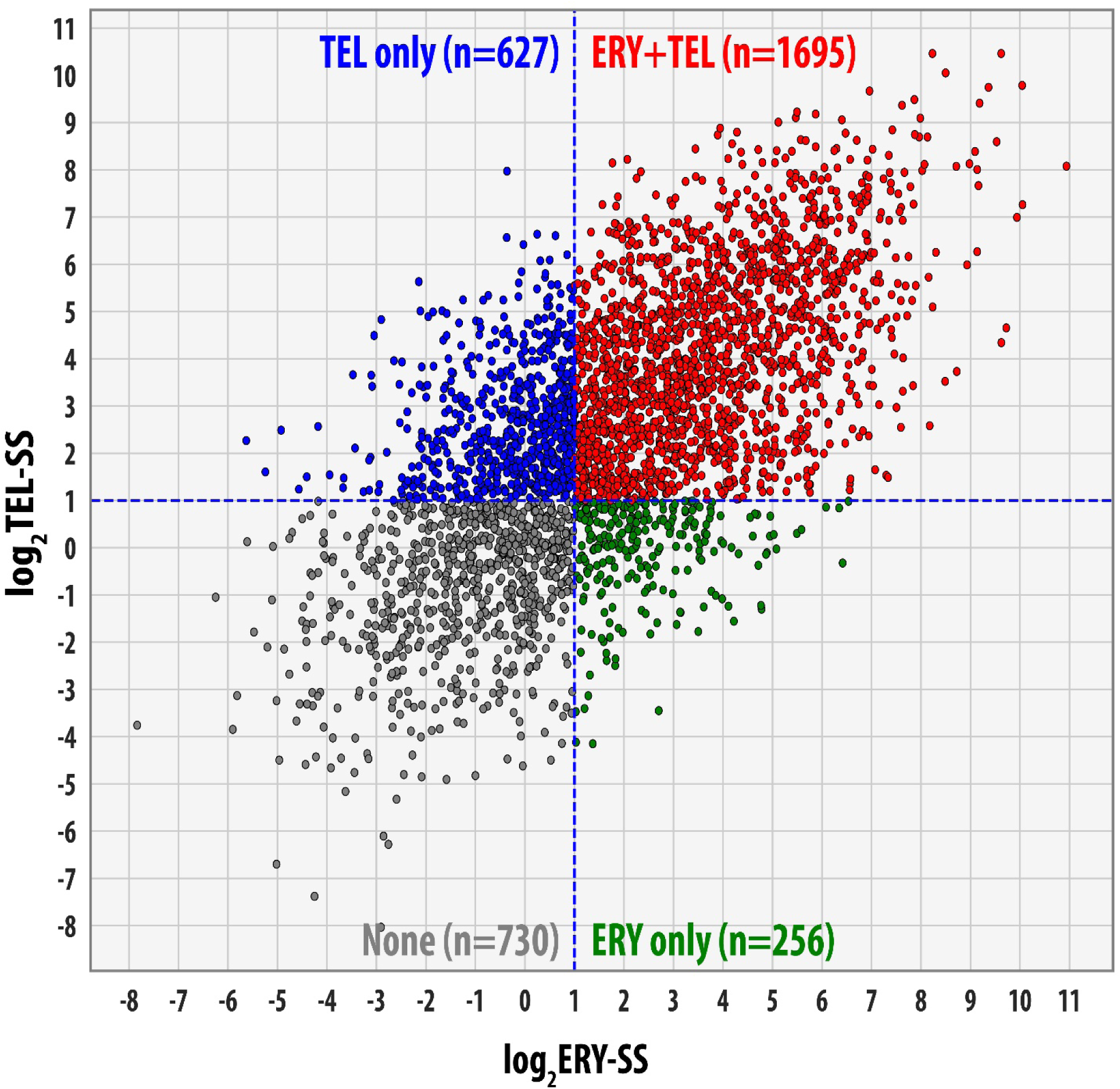
Scatter plot of stalling scores (SS) at +X+ motifs in the presence of ERY and TEL. Ribosome stalling sites are defined as proteome-wide +X+ sites where ribosomal P-site occupancy (position X in the +X+ motif) increases at least twofold in the presence of ERY (X-axis) or TEL (Y-axis) compared to the no-drug control. Dashed blue lines indicate the SS > 2 cutoff values. Proteome-wide +X+ sites unaffected by either macrolide are shown in grey, while ERY- and TEL-specific stalling sites are depicted in green and blue, respectively. Sites where both ERY and TEL induce ribosome stalling are highlighted in red, representing the majority (52%) of all +X+ stalling sites in the *E. coli* proteome.

## SUPPLEMENTARY TABLES

**Table S1.**
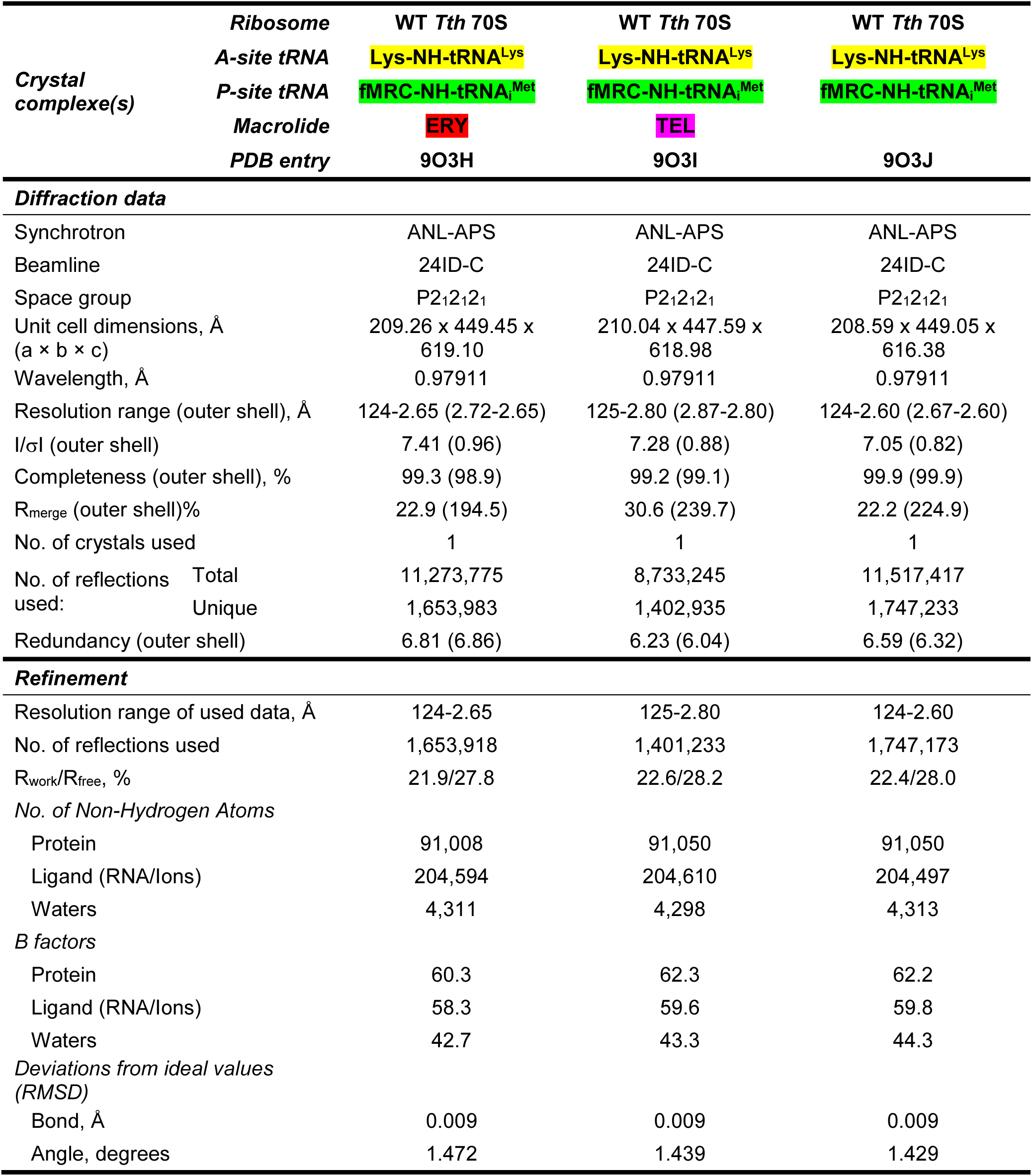
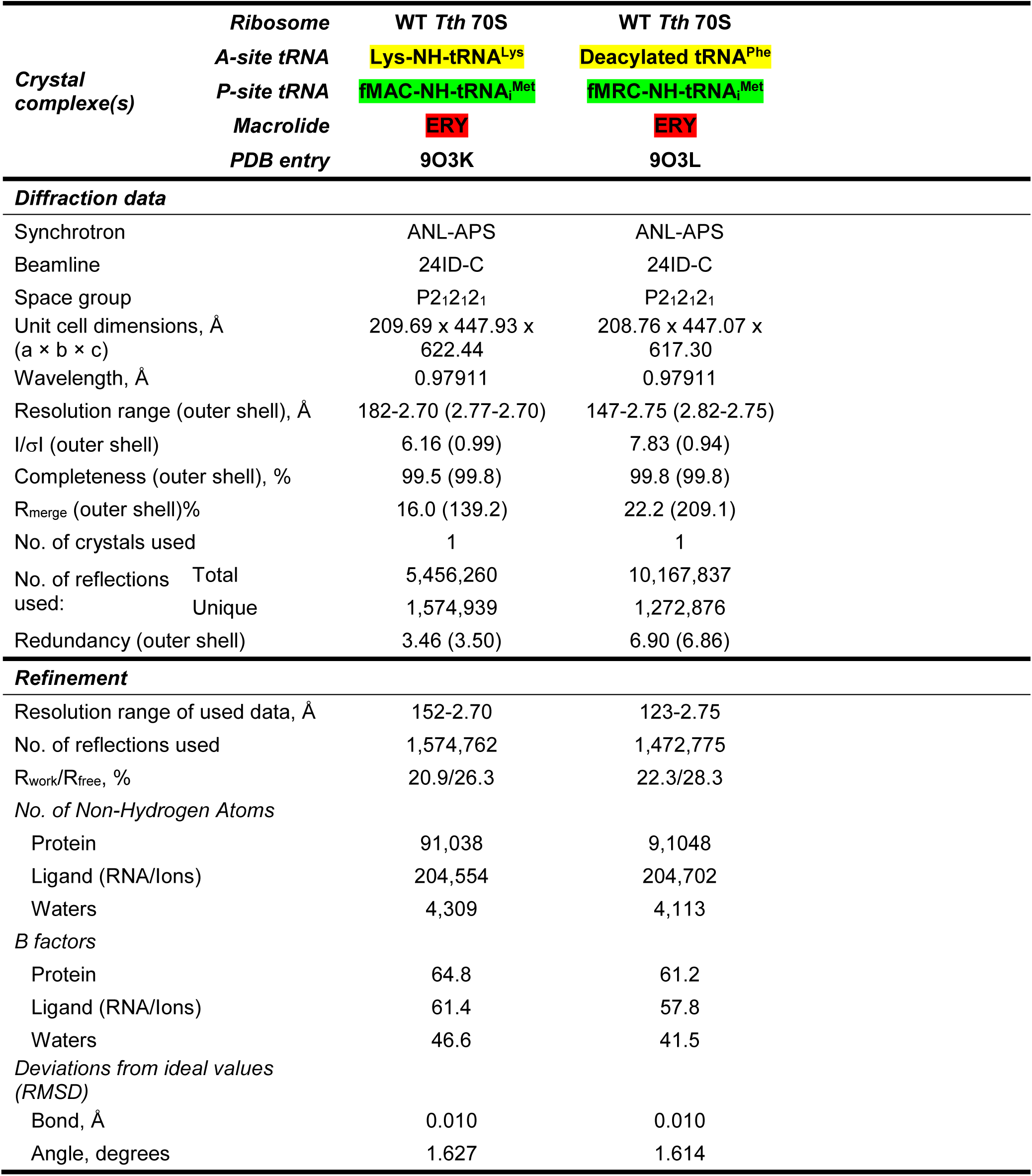
X-ray data collection and refinement statistics.

## MATERIALS AND METHODS

### Reagents

Unless stated otherwise, all other reagents and chemicals were obtained from MilliporeSigma (USA). Macrolides erythromycin and azithromycin were purchased from MilliporeSigma (USA). Ketolide telithromycin was provided by Cempra Inc (USA). All synthetic oligonucleotides, such as DNA primers and mRNAs for structural studies, were obtained from Integrated DNA Technologies (Coralville, IA, USA).

### Toe-printing analysis

The synthetic DNA templates encoding the amino acid sequences for WT and mutant *ermDL* open reading frames (ORFs) were prepared by annealing and extending the long overlapping oligonucleotides listed below (complimentary parts are shown in bold):

- ErmDL-MRLR-Fwd: <u>TAATACGACTCACTATAGGG</u>AGATTTTATAAGGAGGAAAAAATATGCGTCTTCG T**TCGTTCCCAATTACTTGGTAAT**
- ErmDL-MRLK-Fwd: <u>TAATACGACTCACTATAGGG</u>AGATTTTATAAGGAGGAAAAAATATGCGTCTTAAA **TCGTTCCCAATTACTTGGTAAT**
- ErmDL-MRCK-Fwd: <u>TAATACGACTCACTATAGGG</u>AGATTTTATAAGGAGGAAAAAATATGCGTTGCAA A**TCGTTCCCAATTACTTGGTAAT**
- ErmDL-MACK-Fwd: <u>TAATACGACTCACTATAGGG</u>AGATTTTATAAGGAGGAAAAAATATGGCTTGCAA A**TCGTTCCCAATTACTTGGTAAT**
- ErmDL-MRCF-Fwd: <u>TAATACGACTCACTATAGGG</u>AGATTTTATAAGGAGGAAAAAATATGCGTTGCTT C**TCGTTCCCAATTACTTGGTAAT**
- ErmDL-MRCP-Fwd: <u>TAATACGACTCACTATAGGG</u>AGATTTTATAAGGAGGAAAAAATATGCGTTGCCC G**TCGTTCCCAATTACTTGGTAAT**
- ErmDL-Rev: <u>GGTTATAATGAATTTTGCTTATTAAC</u>GACCAAGACAACACTTTTCTAAAT**ATTAC CAAGTAATTGGGAACGA**

The resulting DNA products were amplified by polymerase chain reaction (PCR) using Q5 Hot Stard DNA Polymerase (New England Biolabs, USA) and T7/NV1 primer pair for all variants of *ermDL* sequences:

- T7 primer: TAATACGACTCACTATAGGG
- NV1: GGTTATAATGAATTTTGCTTATTAAC

The sequences of PCR primers within the longer oligonucleotides are underlined.

The toe-printing analysis of drug-dependent ribosome stalling on WT and mutant *ermDL* templates (**Figure S2**) was carried out as previously described^13,32,42–45^ with minor modifications. Toe-printing reactions were carried out in 5-µL aliquots containing a PURExpress transcription-translation coupled system (New England Biolabs, USA) to which the test DNA templates were added. The reactions were incubated at 37°C for 10 minutes. Reverse transcription on the templates was carried out using radioactively labeled NV1 oligonucleotide for 5 minutes at 37°C. Primer extension products were resolved on 6% sequencing gels and, after exposure of the dried gels to the phosphorimager screen, visualized on a Typhoon Biomolecular Imager (Cytiva) as described previously^42^. The final concentration of ERY in the toe-printing was 50 µM. In all reactions, we used 50 µM mupirocin (inhibitor of isoleucyl-tRNA synthetase) to arrest ribosomes at the Ile codon downstream of the macrolide arrest site.

### Preparation of full-length lysine-specific tRNA^Lys^

DNA sequence encoding for lysine-specific tRNA^Lys^ from *E. coli* was cloned into the pBSTNAV plasmid vector (AddGene, USA) using *Eco*RI and *Pst*I restriction sites.

Transformants carrying the resulting pBSTNAV-Lys vector and expressing tRNA^Lys^ were selected by aminoacylation of the total tRNA preparations (50 ng/µL) with 10 µM [^14^C]-lysine (222 dpm/pmol) and lysine-specific aminoacyl-tRNA-synthetase (LysRS) from *Thermus thermophilus* (60 ng/µL) in a buffer containing 60 mM HEPES-KOH pH 7.5, 10 mM MgCl2, 2.5 mM ATP, 5 mM DTT, 0.5 µg/µL BSA. For large-scale preparation of tRNA^Lys^, tRNA-expressing *E. coli* cells were grown overnight in LB media with 100 µg/µL ampicillin. The cells were harvested and resuspended in buffer containing 1 mM Tris-HCl pH 7.4, 10 mM Mg (CH3COO)2, and treated with 0.5 volumes of acidic phenol pH 4.5. The aqueous phase was then precipitated by the addition of two volumes of ethanol and NaCl to 500 mM final concentration. The resulting pellet was resuspended in 1M NaCl, and the soluble fraction, containing total tRNA, was precipitated with ethanol again. The pellet was resuspended in 200 mM Tris-HCl pH 9.0, and the resulting solution of total tRNA was incubated for 2 hours at 37°C to promote deacylation of any remaining aa-tRNAs. Deacylated total tRNA was precipitated with ethanol, resuspended in buffer A (40 mM sodium phosphate buffer pH 7.0), and subjected to anion exchange chromatography on an 8-ml MonoQ column (10/100 GL, GE Healthcare) using 300-ml 0-100% linear gradient of buffer B (40 mM sodium phosphate buffer pH 7.0, 1M NaCl) for elution. Fractions containing tRNA^Lys^ were identified by aminoacylation with [^14^C]-lysine, pooled together, precipitated with ethanol, resuspended in buffer C (20 mM NH4CH3COO pH 5.5, 400 mM NaCl, 10 mM MgCl2, 1 mM EDTA, 1% methanol), and subjected to reversed-phase chromatography on 20-ml C5 column (C5-5, 250×10 mM, Discovery BIO Wide Pore, Supelco) using 300-mL 0-60% linear gradient of buffer C supplemented with 40% methanol. Fractions containing tRNA^Lys^ were identified, pooled, and precipitated with ethanol as before. The final tRNA^Lys^ preparation was resuspended in 10 mM NH4CH3COO pH 5.5 and assessed for purity and ability to accept glycine using denaturing PAGE and aminoacylation assay, respectively. The efficiency of aminoacylation of the final tRNA^Lys^ preparation was estimated to be greater than 95%.

### Tailing of tRNA^Lys^

Tailing of tRNA^Lys^ (replacement of the 3’-terminal A76 nucleotide carrying 3’-OH group with the one carrying 3’-NH2) was performed as described previously with minor modifications for other tRNAs^29,30,46^. Briefly, deacylated tRNA^Lys^ (40 µM) was incubated at 37°C for 1 hour in a buffer containing 100 mM glycine-NaOH pH 9.0, 1 mM DTT, 1 mM pyrophosphate, 1 mM 3’-NH2-ATP (Axxora, USA), and 10 µM of the CCA-adding enzyme from *E. coli*. The reaction was terminated by the addition of EDTA to 20 mM, treated with 1:1 phenol-chloroform mixture (pH 8.0), and precipitated with ethanol. The resulting tRNA pellet was dissolved in 20-40 µL of 10 mM NH4CH3COO pH 5.5 and desalted via Sephadex G-25 (MilliporeSigma, USA) spin columns (20-40 µL of tRNA solution per 500 µL of G-25 media).

### Aminoacylation of tailed tRNA^Lys^

Aminoacylation of the tailed 3’-NH2-tRNA^Lys^ with lysine, 2 µM tRNA was incubated at 25°C for 16 hours in a buffer containing 60 mM HEPES-KOH pH 7.2, 15 mM MgCl2, 20 mM KCl, 1 mM DTT, 5 mM ATP, and 1 mM lysine together with 1 µg/µL lysine-specific aminoacyl-tRNA-synthetase (LysRS) from *E. coli*. The aminoacylation reaction was terminated by the addition of EDTA to 30 mM concentration and then treated with phenol and precipitated with ethanol. The pellet was dissolved in a buffer containing 10 mM HEPES-KOH pH 7.5 to a final tRNA concentration of 20 μg/μL and subjected to reversed-phase chromatography on 1.66-ml C4 column (Jupiter 100 × 4.6 mm, 5 µM, 300Å, Phenomenex, USA) using 40-mL 0-60% linear gradient of buffer C supplemented with 40% methanol. Based on the mobility shift on denaturing PAGE with 7M urea, the total yield of non-hydrolyzable aminoacylated Lys-NH-tRNA^Lys^ was >90%. The aliquots were flash-frozen in liquid nitrogen and stored at -80°C until further use in crystallization experiments.

### *X****-*** ray crystallographic structure determination

Wild-type 70S ribosomes from *T. thermophilus* (strain HB8) were prepared as described previously ^29,31,35,47,48^. Synthetic mRNAs with the sequences 5’-GGC-AA<u>G-GAG-G</u>UA-AAA-AUG-AAA/UUC-UAA-3’ containing a strong Shine-Dalgarno sequence (underlined) followed by the methionine (AUG) and either lysine (AAA) or phenylalanine (UUC) codons were obtained from Integrated DNA Technologies (Coralville, IA, USA). Deacylated tRNA^Phe^ and non-hydrolyzable aminoacylated Phe-tRNA^Phe^ was prepared as described previously^33,35^. Stable amide-linked peptidyl-tRNAs, fMRC-tRNAi^Met^ and fMAC-tRNAi^Met^, were prepared as described previously^29,30^.

Both arrested and non-arrested RNCs were formed by programming 5 μM wild-type *T. thermophilus* tightly-coupled 70S ribosomes with 10 μM synthetic mRNA and incubated at 55°C for 10 minutes, followed by the addition of 10 µM fMRC-peptidyl-tRNAi^Met^, or fMAC-peptidyl-tRNAi^Met^, and 20 µM Lys-tRNA^Lys^, or Phe-tRNA^Phe^, or deacylated tRNA^Phe^ for P and A sites, respectively. For ribosome complexes with ERY or TEL, the antibiotics were added to 500 µM or 250 µM final concentration, respectively. All complexes were formed in the buffer containing 5 mM HEPES-KOH (pH 7.6), 50 mM KCl, 10 mM NH4Cl, and 10 mM Mg(CH3COO)2, and then crystallized in the buffer containing 100 mM Tris-HCl (pH 7.6), 2.9% (v/v) PEG-20K, 9-10% (v/v) MPD, 175 mM arginine, 0.5 mM β-mercaptoethanol. Crystals were grown by the vapor diffusion method in sitting drops at 19°C, stabilized and cryo-protected stepwise using a series of buffers with increasing MPD concentrations (25%, 30%, 35%) until reaching the final concentration of 40% (v/v) MPD as described previously^29–31,35,48^. After stabilization and cryo-protection, crystals were flash-frozen using a nitrogen cryo-stream at 80 K (Oxford Cryosystems, UK).

Collection and processing of the X-ray diffraction data, model building, and structure refinement were performed as described in our recent reports^29–31^. Diffraction data were collected at beamlines 24ID-C and 24ID-E of the Advanced Photon Source (Argonne National Laboratory). A complete dataset for each complex was collected using 0.979-Å X-ray irradiation at 100 K from multiple regions of the same crystal, using 0.3-degree oscillations. Raw data were integrated and scaled using the XDS software (version from Jan 10, 2022)^49^. The initial search model was generated by replacing the A-site Phe-tRNA^Phe^ in the previously published structure of *T. thermophilus* 70S ribosome containing aminoacyl- and peptidyl-tRNAs (PDB entry 8CVK^29^) with the Lys-tRNA^Lys^. Initial molecular replacement solutions were refined by rigid-body refinement with the ribosome split into multiple domains, followed by positional and individual B-factor refinement using the PHENIX software (version 1.17)^50^. Non-crystallographic symmetry restraints were applied to four parts of the 30S ribosomal subunit (head, body, spur, and helix 44) and four parts of the 50S subunit (body, L1-stalk, L10-stalk, and C-terminus of the L9 protein). Structural models were built in Coot (version 0.8.2)^51^. The statistics of data collection and refinement are compiled in **Table S1**. All figures showing atomic models were rendered using the PyMol software (www.pymol.org).

### Analysis of ribo-seq data

Proteome-wide analysis of macrolide-induced ribosome stalling on +X+ motis (**Figure S7**) was performed using RNA-seq reads from accession no. GSE61619^18^. Reads were mapped to the *E. coli* reference genome U00096.3 using GALAXY pipeline^52^, and P-site positions were assigned by counting 15 nucleotides from the 3’-end of the trimmed reads. Ribosome densities were normalized to library size and expressed as reads per million (RPM). The stalling score (SS) was defined as the ratio of ribosome density assigned to the P-site codon (position X in the +X+ motif) in the presence of a drug (either ERY or TEL) to the no-drug control across all +X+ motifs in the *E. coli* proteome. The +X+ sites generating at least twofold more reads in the presence of macrolides were considered stalling sites. Data processing and visualization were performed using custom Python scripts and GraphPad Prism.

## DATA AVAILABILITY STATEMENT

Coordinates and structure factors were deposited in the RCSB Protein Data Bank with accession codes:

- **9O3H** for the wild-type *T. thermophilus* 70S ribosome in complex with mRNA, A-site Lys-tRNA^Lys^, P-site fMRC-peptidyl-tRNAi^Met^, and erythromycin;
- **9O3I** for the wild-type *T. thermophilus* 70S ribosome in complex with mRNA, A-site Lys-tRNA^Lys^, P-site fMRC-peptidyl-tRNAi^Met^, and telithromycin;
- **9O3J** for the wild-type *T. thermophilus* 70S ribosome in complex with mRNA, A-site Lys-tRNA^Lys^, and P-site fMRC-peptidyl-tRNAi^Met^;
- **9O3K** for the wild-type *T. thermophilus* 70S ribosome in complex with mRNA, A-site Lys-tRNA^Lys^, P-site fMAC-peptidyl-tRNAi^Met^, and erythromycin;
- **9O3L** for the wild-type *T. thermophilus* 70S ribosome in complex with mRNA, A-site deacylated tRNA^Phe^, P-site fMRC-peptidyl-tRNAi^Met^, and erythromycin;

All previously published structures that were used in this work for structural comparisons were retrieved from the RCSB Protein Data Bank: PDB entries 6XHX, 6XHY, 7NSP, 7NSQ, and 8CVK.

